# Structure and mechanism of Zorya anti-phage defense system

**DOI:** 10.1101/2023.12.18.572097

**Authors:** Haidai Hu, Thomas C.D. Hughes, Philipp F. Popp, Aritz Roa-Eguiara, Freddie J.O. Martin, Nicole R. Rutbeek, Ivo Alexander Hendriks, Leighton J. Payne, Yumeng Yan, Victor Klein de Sousa, Yong Wang, Michael Lund Nielsen, Richard M. Berry, Marc Erhardt, Simon A. Jackson, Nicholas M.I. Taylor

## Abstract

Zorya is a recently identified and widely distributed bacterial immune system, which protects against phage invasion. It consists of a predicted membrane-embedded complex (ZorAB) and soluble components that differ among Zorya subtypes, notably ZorC and ZorD, in type I Zorya systems. Here, we reveal the molecular basis of the Zorya defense system using cryo-electron microscopy, mutagenesis, fluorescence microscopy, proteomics, and functional studies. We demonstrate that ZorAB shares the stoichiometry of other 5:2 inner membrane ion-driven rotary motors. Additionally, ZorA_5_B_2_ features a dimeric ZorB peptidoglycan binding domain and a pentameric α-helical coiled-coil tail made of ZorA that projects approximately 700 Å into the cytoplasm. We further characterize the structure and function of the soluble Zorya components, ZorC and ZorD, and find that they harbour DNA binding and nuclease activity, respectively. Comprehensive functional and mutational analyses demonstrates that all Zorya components work in concert to protect bacterial cells against invading phages. We present evidence that ZorAB operates as an ion-driven motor that becomes activated and anchors to the cell wall upon sensing of cell envelope perturbations during phage invasion. Subsequently, ZorAB transfers the phage invasion signal through the ZorA cytoplasmic tail to the soluble effectors, which function to prevent phage propagation. In summary, our study elucidates the foundational mechanisms of Zorya function and reveals a novel triggering signal for the rapid activation of an anti-phage defense system.

## Introduction

Bacteria and Archaea are frequently attacked by bacteriophages (or phages). Microbes have developed an array of defense strategies to counteract these phage infections^1–3^. Such defense strategies include restriction-modification (R-M) systems that target phage genomic DNA; CRISPR-Cas systems, offering sequence-specific acquired immunity that enables cells to recognize, remember previous and combat future phage infections; abortive infection (Abi) systems that lead to metabolic arrest or cell death upon phage invasion, as well as other defense systems with mechanisms which have only recently begun to be uncovered^4,5^. Anti-phage defense systems not only play a pivotal role in regulating bacterial populations by balancing phage-host dynamics but also present great opportunities for the development of novel biotechnological tools^6^. Recently, through systematic analysis of microbial genomes, several new gene families that protect against phage infections have been identified^7–10^. Further studies have provided key insights into how these defense systems are triggered. These triggering factors commonly include phage structural components, other phage-encoded proteins, or phage nucleic acids that are recognized directly or indirectly by host defense proteins^11,12^. Considering that the initial steps of phage invasion involve interactions with the cell envelope, some defense systems may detect signals originating from the changes in the cell envelope during the early stage of infection. However, such defense mechanisms have so far remained elusive.

Among the most widespread of many recently discovered anti-phage defense systems are the Zorya systems, for which three subtypes have been identified^7,8^. Shared among all Zorya subtypes are two membrane proteins, ZorA and ZorB, which are proposed to form an oligomeric channel complex and contain domains related to the bacterial flagellar stator unit MotAB^7^. Integral to the cytoplasmic membrane, MotAB utilizes the proton-motive force across the inner membrane to drive rotation of the bacterial flagellum^13–16^. In addition to the proposed membrane complex, Zorya defense systems contain one or two cytosolic proteins with limited similarity to other proteins and hitherto unknown functions. These soluble proteins vary among different subtypes, with type I Zorya systems harboring ZorC and ZorD. Displaying limited homology with MotA and MotB, ZorA and ZorB in combination with their respective intracellular Zorya proteins, presumably have evolved and acquired adaptations to counter phage infection.

Of the few characterized bacterial immune systems that contain membrane-associated proteins, most appear to employ defense mechanisms wherein activation of membrane-anchored proteins disrupts or depolarizes the host cell membrane^5,17^. This membrane interference leads to the death or dormancy of infected cells before phages complete their replication cycle, preventing the release of viable progeny, a mechanism typically termed abortive infection^18^. An alternative to abortive infection is direct defense, in which a defense system clears an infection without causing host cell death or dormancy. It has been suggested that ZorAB may function as an ion channel that facilitates membrane depolarization during abortive infection^7^. However, it has not been ruled out that ZorAB could instead act as the sensor of infection, and the molecular details and functional mechanisms of Zorya systems remain to be elucidated.

In this study, we combined single-particle cryogenic electron microscopy (cryo-EM), mutagenesis, functional assays, proteomics, and total internal reflection fluorescence microscopy to decipher several key aspects of the Zorya defense mechanism. We discovered that ZorA and ZorB form a unique 5:2 ion-driven motor complex in the inner membrane that contains a long, intracellular tail structure. We provide evidence that Zorya is a direct defense system, where ZorAB is a sensor responsible for detecting phage-induced perturbations of the cell envelope. During sensing of phage invasion, we propose that the dimeric ZorB peptidoglycan-binding domain anchors the ZorAB complex to the cell wall near the phage injection site and that the ion-driven motor rotates the ZorA tail within the cytosol. The rotation would then transmit the invasion signal to the effector subunits ZorC and ZorD, with ZorD being recruited to the site of activated ZorAB. ZorC and ZorD display *in vitro* DNA binding and autoinhibited nuclease activity, respectively. Together, activated ZorC and ZorD target and degrade the invading phage DNA within the proximity of the ZorA tail, thereby preventing phage propagation. A long rotating tail-spike located close to the site of phage DNA injection is strongly suggestive of a “bobbin” around which the invading DNA could be wound, thereby immobilizing it for subsequent degradation.

## Results

### Zorya has broad activity against phage invasion through a direct immunity mechanism

Type I and II Zorya systems are widely distributed across Gram-negative phyla, whereas type III Zorya appears to primarily occur within *Pseudomonadota* (**Fig. 1a**). Intriguingly, Zorya is rarely found in Gram-positive bacteria. To better understand the Zorya anti-phage defense mechanism, we investigated the activity of an *E. coli* type I Zorya system (*Ec*ZorI). We cloned the complete *Ec*ZorI operon from strain NCTC9026, including its native promoter, into a low copy plasmid (pACYC) and used a heterologous *E. coli* strain (MG1655 ΔRM) to examine *Ec*ZorI-mediated anti-phage defense against a panel of 70 phages^19^. The *Ec*ZorI system provided anti-phage activity against a diverse range of phages, including different families (**Fig. 1b**). Phage adsorption was not affected by *Ec*ZorI, indicating that anti-phage defense occurs at a subsequent stage of infection (**Fig. 1c**). *Ec*ZorI also prevented the phage burst of infected cells (**Fig. 1d**). We next examined whether *Ec*ZorI affected incoming DNA during bacterial conjugation or plasmid transformation but found no evidence of *Ec*ZorI defending against these mechanisms of foreign DNA uptake (**Extended Data Fig. 1a, b**). Therefore, our findings indicate that some aspect of phage infection, other than the mere introduction of foreign DNA into the cell, is required to trigger Zorya activity.

**Figure 1.**
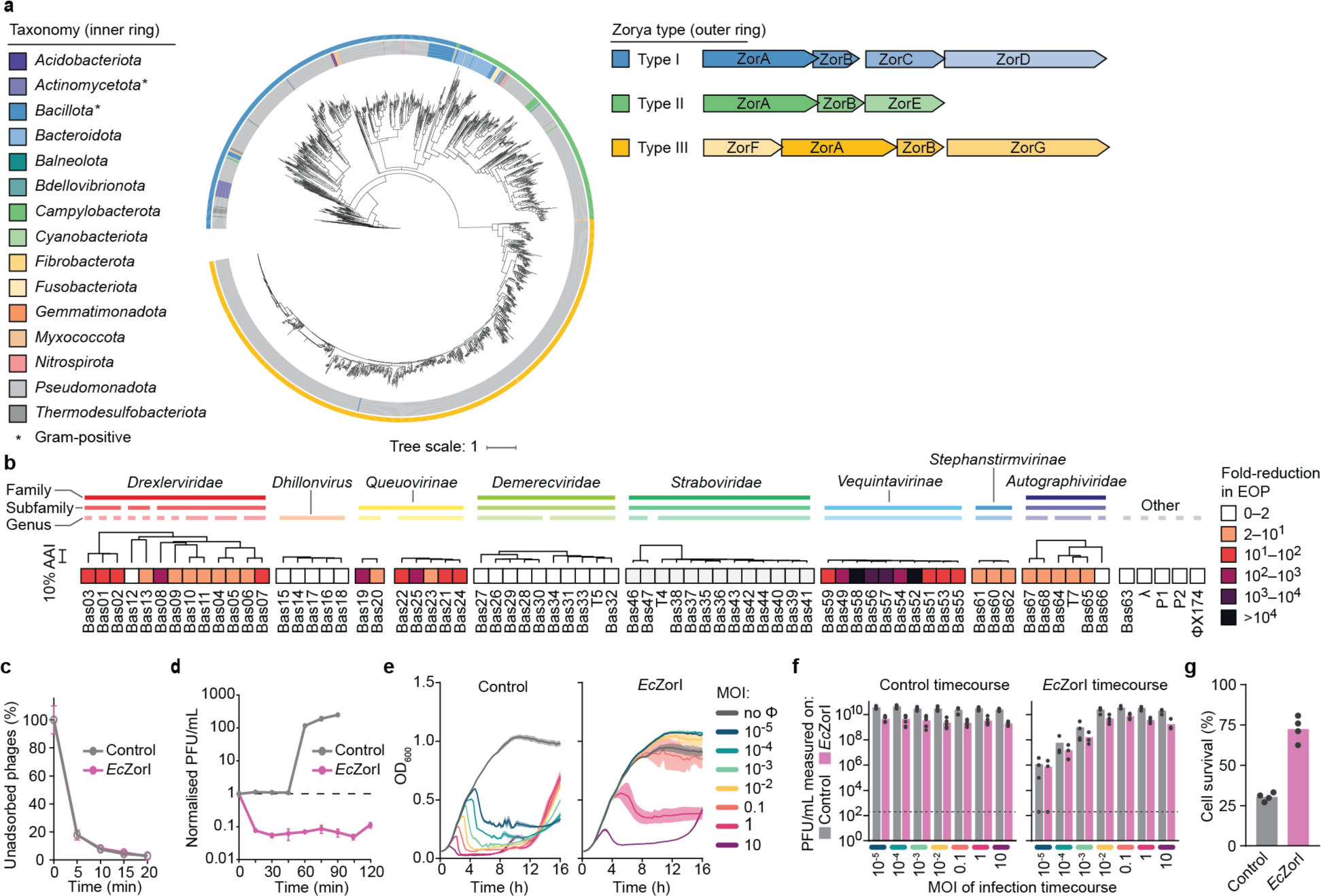
Zorya has broad activity against phages via a direct immunity mechanism. **a**, Zorya phylogenetic tree. As ZorA and ZorB are present in all Zorya types, the tree was generated using concatenated ZorA+B sequences. The outer ring represents the Zorya types I, II or III, and the inner ring represents the taxonomy of bacteria. The taxonomic rank being highlighted is at the phylum level. On the right, the gene organization of the three Zorya types is shown. **b**, *Ec*ZorI defense against diverse *E. coli* phages, determined using efficiency of plaquing (EOP) assays. AAI: Average amino acid identity. **c**, Adsorption of phage Bas24 to *E. coli* cells possessing or lacking *Ec*ZorI. **d**, One-step phage growth curve for phage Bas24 infecting *E.coli*, with or without *Ec*ZorI, normalized to the plaque forming units (PFU) per mL at the initial timepoint. **e**, Infection time courses for liquid cultures of *E. coli*, with and without *Ec*ZorI, infected at different multiplicities of infection (MOI) of phage Bas24. **f**, Phage titers at the end timepoint for each sample from the infection time courses (e), measured as EOP on indicator lawns of *E. coli* either without (control) or with *Ec*ZorI. The limit of detection (LOD) is shown with dotted lines. **g**, Survival of *E. coli* cells, lacking or possessing *Ec*ZorI, infected at an MOI of 5 with Bas24. In panels **b-d**, data represent the mean of at least three replicates and error bars (**c,d**) or shaded regions (**e**) represent the standard error of the mean (SEM).

Next, we examined whether the *Ec*ZorI system provided population-level defense in liquid cultures infected at different multiplicities of infection (MOI) (**Fig. 1e** and **Extended Data Fig. 1c**). Although each phage tested affected the control populations to differing extents, population growth in the *Ec*ZorI samples was generally unaffected at low (<0.1) MOI and in some cases also high (>1) MOI. Importantly, the growth kinetics at early timepoints did not reveal any premature host population collapse or delayed growth for cells expressing *Ec*ZorI compared to the empty vector controls, as might be expected for abortive defense mechanisms. Phages were detectable at the end timepoints in most infected cultures, but at lower levels in the presence of *Ec*ZorI (**Fig. 1f and Extended Data Fig. 1d**). These findings suggest the *Ec*ZorI system does not act via an abortive infection mechanism, where induced cell dormancy or death of infected cells leads to phage extinction and population-level protection. To test this further, we measured the survival rate of cells infected with Bas24 and found that *Ec*ZorI increased cell survival, again consistent with a direct defense mechanism rather than abortive infection (**Fig. 1g**).

### Zorya contains a ZorA_5_B_2_ complex with a unique intracellular tail

To elucidate the molecular assembly of ZorA and ZorB, we determined the structure of the *Ec*ZorI ZorA and ZorB complex (*Ec*ZorAB) using single-particle cryo-EM. We expressed then purified the *Ec*ZorAB complex from cell membranes using the mild detergent lauryl maltose neopentyl glycol (LMNG) (**Fig. 2a** and **Extended Data Fig. 2a**). Visualized by negative stain EM, *Ec*ZorAB appeared to contain a ‘head’ domain attached to a long tail-like structure measuring approximately 700 Å (**Fig. 2b**). We then resolved the *Ec*ZorAB cryo-EM structure to an overall resolution of 2.7 Å (**Extended Data Fig. 2b-c** and **Extended Data Table 1**). The EM reconstruction reveals an oligomeric complex consisting of five ZorA subunits and two ZorB subunits, with the same stoichiometry and arrangement in the membrane as the flagellar stator unit MotA–MotB complex (MotA_5_B_2_)^15^ (**Fig. 2c-e, h**). Quantitative mass spectrometry analyses of *Ec*ZorI-expressing *E. coli* cells support the 5:2 ZorA:ZorB ratio (**Extended Data Fig. 2k, l** and **Extended Data Table 2**). Overall, *Ec*ZorAB comprises four distinct structural layers: a peptidoglycan binding domain (PGBD, ZorB residues T47-L246), transmembrane domain (TMD), membrane-proximal cytoplasmic domain (MPCD), a region spanning ZorA residues G48-L127, K207-S222), and a tail-like structure formed by the ZorA C-terminal region (ZorA residues G223-T729) (**Fig. 2a**, **2d**).

**Figure 2.**
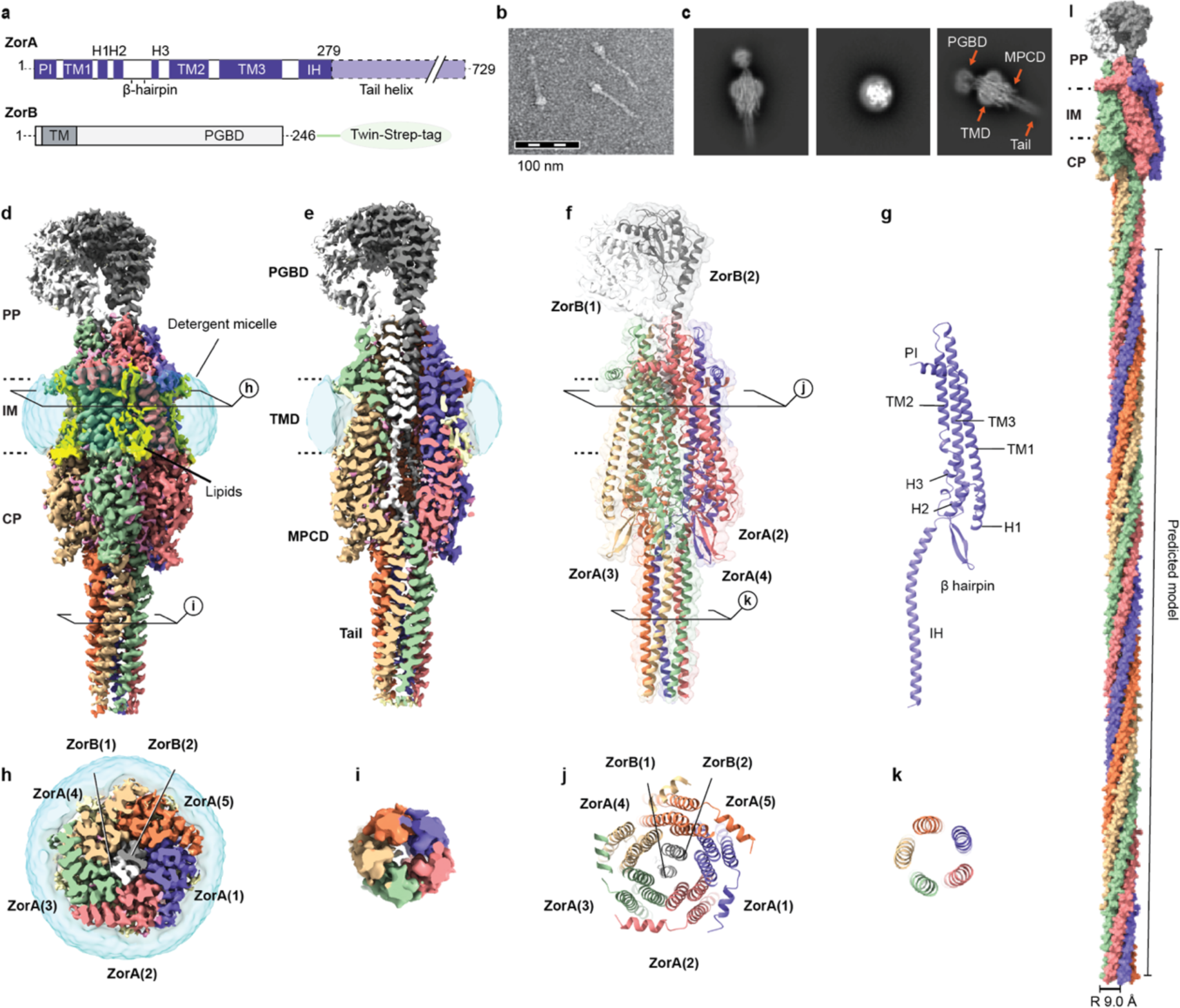
Cryo-EM of *Ec*ZorAB and its architecture. **a**, Schematic representation of *Ec*ZorA and *Ec*ZorB. **b**, Negative staining image of purified *Ec*ZorAB particles. **c**, Representatives of high-resolution 2-dimensional classes of *Ec*ZorAB particles from cryo-EM. Domain architectures of the E*c*ZorAB complex are depicted. **d**, Cryo-EM map of *Ec*ZorAB. Five ZorA subunits (purple, salmon, light green, tan, and coral) surround two ZorB subunits (white and dark gray) viewed from the plane of the membrane. Membrane-bound lipids are shown in yellow. The detergent micelle is shown as a translucent surface representation in cyan. Dashed lines depict inner membrane boundaries. **e**, Cross-section view of the EM density map. **f**, Ribbon model representation of *Ec*ZorAB. **g**, Structure of a single ZorA subunit. **h**-**i**, Cross-section view of the Cryo-EM map of *Ec*ZorAB TMD (**h**) and tail (**i**) from the periplasmic side. **j**-**k**, Cross-section view of the model of *Ec*ZorAB TMD (**j**) and tail (**k**) from the periplasmic side. **l**, Composite model of *Ec*ZorAB whole complex. the predicted portion of the ZorA tail is represented as surface. The radius of the ZorA tail is shown in **l**. PP, periplasm; IM, inner membrane; CP, cytoplasm; PGBD, peptidoglycan binding domain; TMD, transmembrane domain; MPCD, membrane-proximal cytoplasmic domain; TM, transmembrane; H, helix.

The periplasmic region of the complex exhibits flexibility relative to the TMD. Nevertheless, local refinement improved the resolution to 3.5 Å, clearly resolving a dimerized ZorB PGBD (**Extended Data Fig. 2h-j**). The resolution corresponding to the TMD and MPCD reached 2.2 Å, allowing us to model sidechain rotamers and non-protein molecules such as water, ions and lipids. For the ZorA tail, our cryo-EM map provides density information for the first 56 residues. Mass spectrometry analyses on the purified *Ec*ZorAB complex confirmed the presence of the intact ZorA C-terminal region, consistent with the negative stain images, but conformational heterogeneity prevented 3D reconstruction of the entire ZorA tail (**Fig. 2b-d** and **Extended Data Fig. 3a**). Secondary structure prediction revealed a preference for the tail to adopt α-helical structures, suggesting that the rest of the ZorA tail is likely to be a continuation of the experimentally determined structure, which folds into a coiled coil with a right-handed super-helical twist towards cytoplasm (**Extended Data Fig. 3a**). Additionally, we observed a consistent hydrophobic pattern in the tail sequence (**Extended Data Fig. 3c, d)**. Based on these observations, we constructed an idealized full-length ZorA model where the ZorA tail forms a helical bundle projecting into the cytoplasm, having a helical pitch of 328 Å and a radius of 9.0 Å (**Fig. 2l** and **Extended Data Fig. 3c**).

### ZorAB is a peptidoglycan-binding rotary motor

On the periplasmic side, the C-terminal PGBDs of the two ZorB subunits form a homodimer, with each monomer consisting of four helices (α1-α4) and an antiparallel β-sheet (β1-β5) (**Fig. 3a** and **Extended Data Fig. 4a, b**). The dimerization interface is composed of α3 and β5 from each monomer, driven mainly by van der Waals forces and electrostatic interactions. Additionally, a C-terminal loop from ZorB caps the side of the dimerization interface (**Fig. 3b**). Each monomer contains two disulfide bridges resolved in the cryo-EM density: one connects the α1 and β1-β2 loops, and the other connects α3 and the ZorB C-terminal end, potentially contributing to the folding, stability and rigidity of the ZorB PGBDs (**Fig. 3b** and **Extended Data Fig. 4c, d**). The overall ZorB dimer structure resembles that of the periplasmic domain of the flagellar stator units MotB, and other peptidoglycan (PG) binding proteins^20,21^ (**Extended Data Fig. 4b**). MotAB is kept in an inactive state by the MotB ‘plug’ regions (following the TMD and preceding the PGBD) that inhibit ion flux and rotation of MotA around MotB (**Extended Data Fig. 4f, g**). Only upon incorporation of MotAB into the flagellar motor is the MotB plug released and the PGBDs dimerize to enable PG binding^15^. Intriguingly, in our structure the ZorB PGBD is already dimerized and purified ZorAB can bind PG (**Extended Data Fig. 4e**).

**Figure 3.**
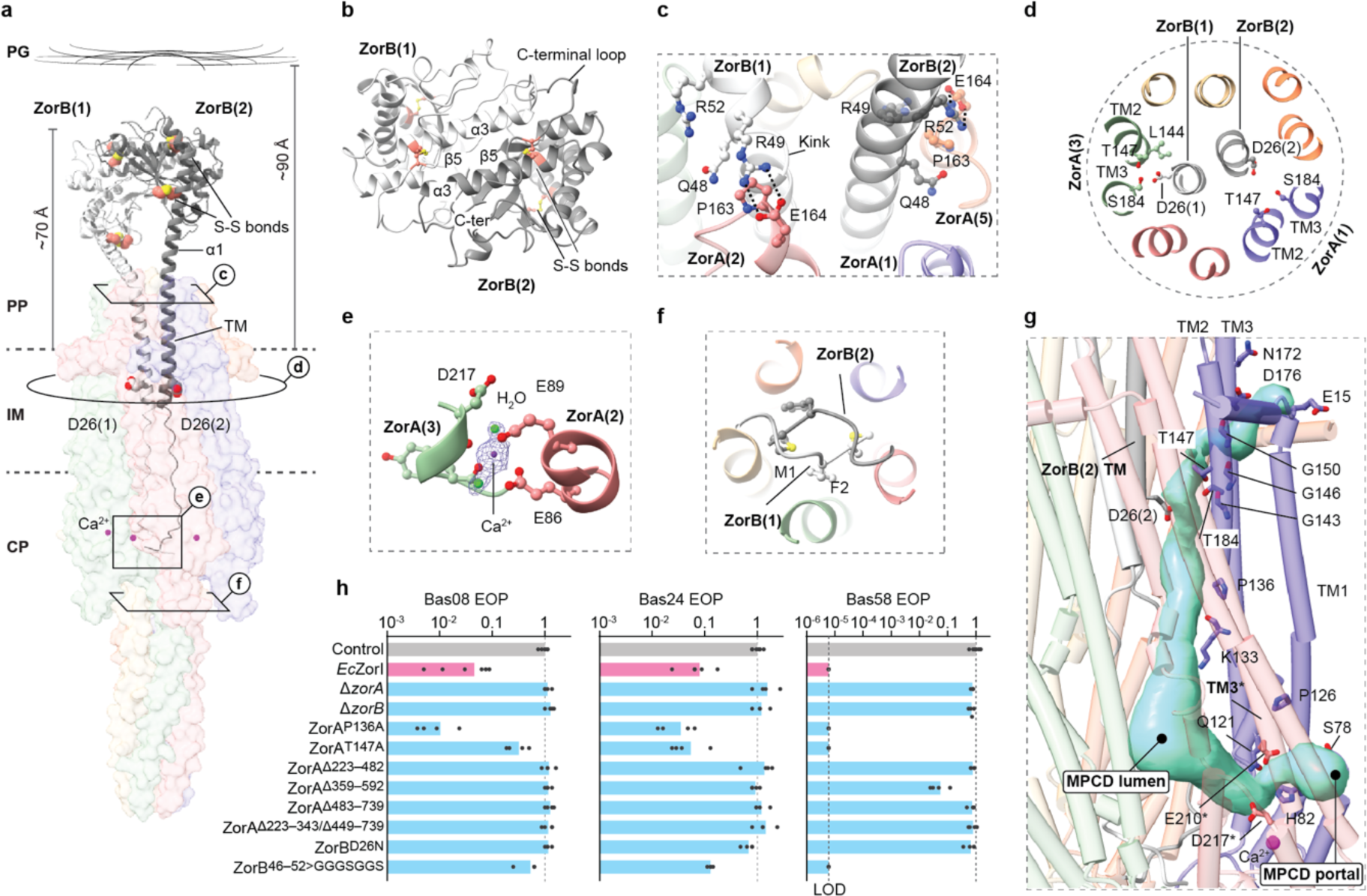
ZorAB is a PG-binding rotary motor. **a**, *Ec*ZorAB viewed from the plane of the membrane, with ZorB shown as ribbons (black and white) and ZorA shown as a translucent surface representation. The distance between the inner membrane and PG layer in *E.coli* is approximately 90 Å^68^. The cysteines from the two disulfide bridges in the ZorB PGBD are indicated and shown as spheres. The aspartate residues D26 from both ZorB TM are indicated and shown. **b**, Top view of the ZorB PGBD. **c**, Close-up view of the interactions of ZorB with ZorA at the domain assembly interface in the periplasmic space. **d**, Cross section view of ZorAB TMD, showing the ZorB D26 and surrounding residues. **e**, Ca^2+^ binding site. EM densities are only overlapped on Ca^2+^ ion, and the two water molecules. **f**, Close-up view of the interactions of the ZorB N-terminus with ZorA tail **g**, Ion translocation pathway (semitransparent surface representation in light blue) in ZorAB. Residues along the ion permeation pathway and from the ion selectivity filter are shown. Each asterisk indicates residues or structural elements from neighboring ZorA subunit. **h**, The effects of ZorA and ZorB mutations on *Ec*ZorI-mediated anti-phage defense, as measured using EOP assays. Data represent the mean of at least 3 replicates and are normalized to the control samples lacking *Ec*ZorI. For phage Bas58, the limit of detection (LOD; one plaque observed in the assay) is marked with a bashed line. In the absence of any observed plaques, the result was recorded as the LOD. ZorB^46–52>GGGSGGS^ corresponds to the replacement of ZorB residues 46-52 with a GGGSGGS linker. Data for additional phages are provided in **Extended Data Fig. 6a**.

In the ZorAB TMD, two ZorB transmembrane helices (TM) are asymmetrically surrounded by five ZorA subunits. Each ZorA subunit consists of three transmembrane helices (TM1-TM3). ZorA TM2 and TM3 are lined directly against ZorB TM, while TM1 is peripheral and faces the lipid bilayer (**Fig. 2g**). Lipid densities are observed in the EM reconstruction on both leaflets of the membrane (**Extended Data Fig. 2f, g**). These protein-bound lipids likely stabilize the TMD and enhance the oligomeric assembly. The TMD of ZorAB is structurally related to that of 5:2 ion-driven prokaryotic rotary motors, including possessing the universally conserved and mechanistically important aspartate residue, D26 in ZorB. One ZorB D26 is engaged with ZorA chain C TM2 T147 and TM3 S184 and the other D26 is unengaged and points toward a lumen enclosed by the ZorA MPCD (**Fig. 3a, d**). The interaction modes of these two D26 are the same as those in the inactive state of MotAB, suggesting similar working mechanisms (**Extended Data Fig. 4h-j**). Additionally, we do not find any region corresponding to a ‘plug’ in ZorB. However, we do observe that P163 from ZorA(2) (ZorA chain 2) induces a kink in ZorB(1) α1 near residue 46, and two pairs of salt bridges (ZorA(2) E164-ZorB(1) R49; ZorA(5) E164-ZorB(2) R52) and several polar interactions are located at the ZorAB periplasmic assembly interface that might block the rotation of ZorA around ZorB in this conformational state (**Fig. 3a, c**). In support of a rotation-suppressing role, when we replaced ZorB residues 46-52 with a GGGSGGS linker we were able to generate a nearly identical Cryo-EM reconstruction to the WT complex, apart from the unresolved density of ZorB TMDs and the flexible ZorB PGBDs (**Extended Data Fig. 4m-q**), which is consistent with freedom of ZorA to rotate around ZorB (through Brownian motion) in the mutant.

On the cytoplasmic side, the ZorA TMD and MPCD are directly connected by TM1 and TM3, the intracellular segments of which are joined by three vertical helices (H1-H3) and a β-hairpin motif (**Fig. 2g**). H3 is less ordered due to the presence of two proline residues, P126 and P136. We found five strong, spherical densities in the ZorA MPCD, each coordinated by the mainchain carboxylate groups of S218 and Y220 at the end of TM3, and the side chains of E86 and E87 from the adjacent subunit, as well as two well-resolved water molecules (**Fig. 3a, e**). Based on the strongly negative electrostatic environment and the surrounding coordinating residues, we assigned this density to a Ca^2+^, which bridges the MPCD of two adjacent ZorA subunits and links ZorA TM3 to its intracellular helix (**Fig. 3e** and **Extended Data Fig. 5b**).

Consistent with a function for ZorAB as an ion-driven rotary motor, we observed a water-filled ion permeation pathway connecting the periplasmic space, via the unengaged ZorB D26, to the cell cytosol (**Fig. 3g**). On the periplasmic side, a cavity is lined by the end of TM2, the beginning of TM3, and the turn between the ZorA periplasmic interface (PI) helix and TM1. Several negatively charged residues are found in this region, which join to create a negatively charged environment that would attract incoming ions (**Fig. 3g**). Moving towards the cytoplasmic side, ZorA residues T147 and S184 resemble an ion selectivity filter^16^ that controls ion access from the periplasm to ZorB D26 (**Fig. 3d, g**). The absence of the additional polar residues in the ion selectivity filter strictly required for sodium coordination indicates that ZorAB is likely a proton-driven motor^16^ (**Extended Data Fig. 4h-j**). The pathway extends from ZorB D26 in the direction of the cytoplasm to the inner lumen encircled by the ZorA MPCD. We found lateral portals in the ZorA MPCD, framed on one side by H2 and the broken H3, and on the other side, by the intracellular part of TM3 from the neighboring subunit. The portal is beneath the head of the membrane-bound lipids and above the Ca^2+^-binding site, and it is highly hydrated, which could facilitate ion exit (**Fig. 3g**). Taken together, our structural analyses and comparisons imply that upon activation, the ZorAB TMD utilizes the across membrane proton gradient to drive the rotation of ZorA around ZorB.

Next, we mutated residues along the ion-permeation pathway to assess the role of the ZorAB TMD in Zorya anti-phage defense activity. ZorB D26 is universally conserved and essential for all models of ion translocation and motor rotation. Replacement with asparagine abolishes defense against phage infection (**Fig. 3d, h** and **Extended Data Fig. 6a**). In the ion-selectivity filter, mutation of ZorA T147 to alanine significantly decreases defense ability. Furthermore, mutation of ZorA P136, which creates a kink in the ZorA MPCD H3 helix and in turn leads to the formation of an electrostatic contact between the backbone carbonyl oxygen of ZorA(4) K133 and the side chain of ZorB(1) K18, to alanine results in increased defense activity (**Fig. 3g, h** and **Extended Data Fig. 6a**). These observations are consistent with rotation of ZorAB, driven by ion flux through the above pathway, being essential for Zorya anti-phage defense.

### ZorAB intracellular tail controls the Zorya anti-phage defense activity

One of the most salient features of the ZorAB complex is its long tail-like structure, of which we could confidently model the initial 56 residues (residues G223-T279) (**Fig. 2a, d**). Within the ZorA MPCD, ZorB N-terminal residues M1 and F2 intertwine and hydrophobically block the entrance of the tail (**Fig. 3f**). On the outside of the tail, residue R108 from the β-hairpin motif forms a salt bridge with D22, and H92 makes electrostatic contact with the hydroxyl group of F228 (**Extended Data Fig. 5a, b**). These interactions appear to be critical for the assembly of the ZorAB complex, since disrupting these interactions by deleting the entire tail (residues G223-T729) abolish ZorAB complex formation, and we could only purify the dimerized ZorB (**Extended Data Fig. 5e**). Inside the tail there are many hydrophobic residues present. L250, L254, L258 and L261 from each ZorA subunit comprise continuous hydrophobic pentameric rings. Additionally, we observe an extra density along the tail central axis in this region, which is best modeled as a fatty acid, consistent with a predicted lipid binding site by deep learning methods^22^ (**Extended Data Fig. 5c**). Given that the tail structure protrudes into the cytoplasm and is surrounded by aqueous solution, hydrophobic interactions inside the tail seem to be the primary driving force for tail assembly, and it is unlikely that the tail conducts ions or other small soluble molecules. Intriguingly, part of the ZorA tail (residues 540-729) shows homology with the core signaling unit of the bacterial chemosensory array (**Extended Data Fig. 3b**), which contains a long intracellular helical bundle responsible for transferring signal from the extracellular environment into the cell and regulates the activities of the subsequent effectors^23^. Sequence analyses further reveal that the length of the tail appears consistent across Zorya subtypes, suggesting that a uniform tail length is functionally essential (**Extended Data Fig. 6b**).

Deletion of any of the Zorya proteins results in loss of anti-phage protection, emphasizing that the complete function of the Zorya system requires the presence of all its components and relies on the communication between the membrane anchored ZorAB complex and cytosolic soluble proteins (**Fig. 3h** and **Fig. 4c, h**). Considering the motor-like structural features of ZorAB TMD and its intracellular long tail, we speculated that the ZorA tail is responsible for transmitting a signal derived from the motor activity to the cytosolic proteins ZorC and ZorD. We assessed whether the integrity of the ZorA tail is required for the system’s function. We made four ZorA tail truncations: deleting the beginning (ZorA^Δ223–482^), middle (ZorA^Δ359–592^) and tip (ZorA^Δ483–729^) of the tail as well as a combination of deleting the beginning and middle (ZorA^Δ223–343/Δ449–729^). All these mutations abolish *Ec*ZorI anti-phage defense ability (**Fig 3h**, **Extended Data Fig. 5a** and **Extended Data Fig. 6a**). To discern whether deletion of the middle or tip of the tail affect ZorAB assembly, we further expressed and determined the structures of two ZorAB mutants (ZorA^Δ359–592^ and ZorA^Δ435–729^). Examination of the purified mutant samples under negative stain EM show the shortened tail length (**Extended Data Fig. 5f, g**). Cryo-EM structures reveal that the two mutants have similar assemblies to ZorAB WT, unlike the complete tail deletion. (**Extended Data Fig. 5h, i** and **Extended Data Table 1**). Hence, truncation of the middle or the tip of the tail does not affect the ZorAB TMD motor assembly or formation of the remainder of the tail bundle but does impede Zorya function. Subsequently, the Ca^2+^ binding site was also investigated. Indeed, double mutation of E86 and E89 to alanine results in loss of Zorya function (**Fig. 3h, Extended Data Fig. 5j** and **Extended Data Fig. 6a**). Structural comparisons between the Ca^2+^ mutant and ZorAB WT reveal conformational changes in the ZorA MPCD, including the linker between TM3 and ZorA tail helix (**Extended Data Fig. 5d, j**). Inactive Ca^2+^ binding sites therefore likely break the connection between the ZorAB TMD motor and the tail. Collectively, these results demonstrate that ZorAB tail length and integrity are essential for Zorya protection from phage infection.

**Figure 4.**
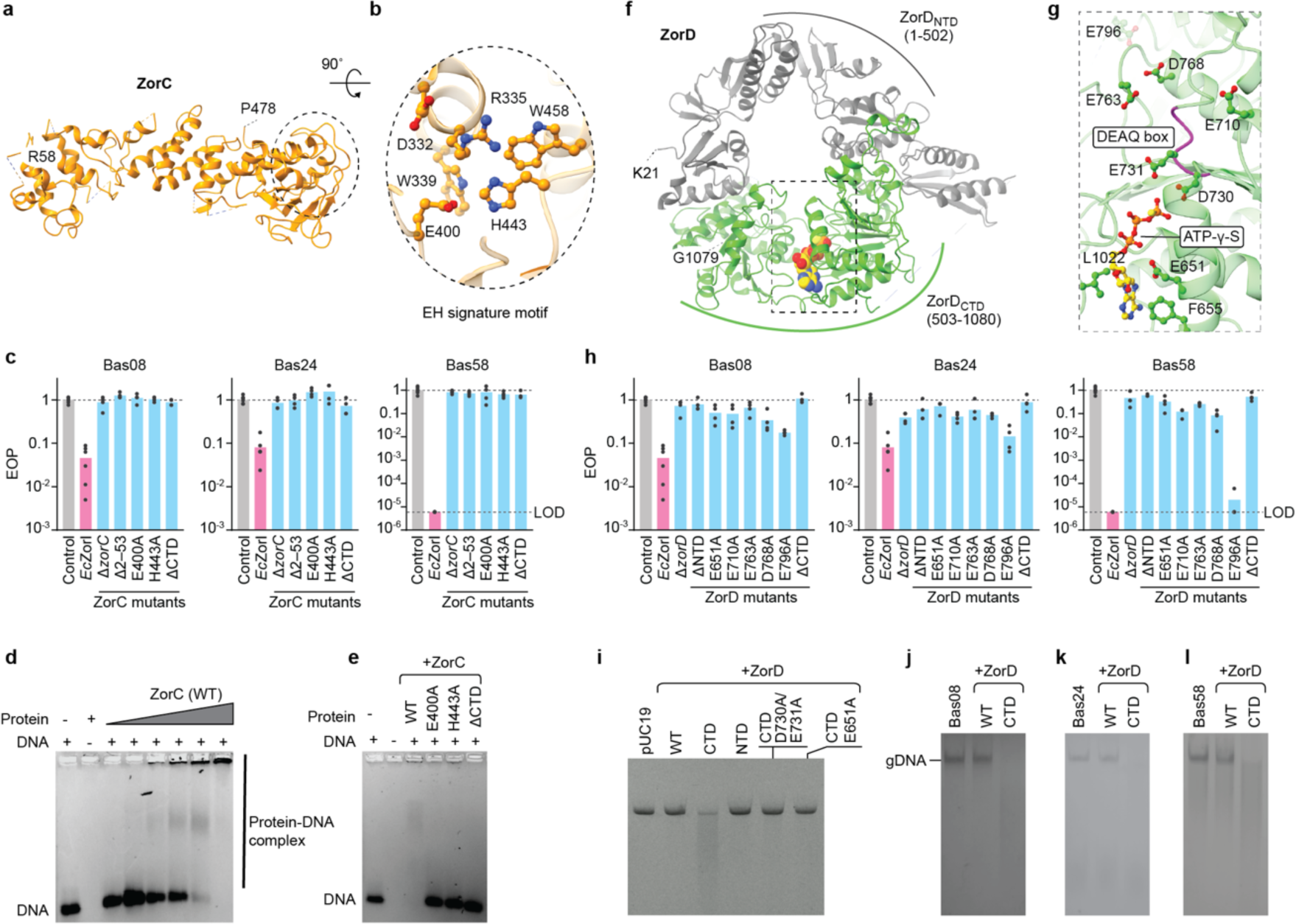
Structural and functional characterization of ZorC and ZorD. **a**, Ribbon model representation of ZorC. **b**, Details of the ZorC EH signature motif. **c**, The effects of ZorC mutations on *Ec*ZorI-mediated anti-phage defense, as measured using EOP assays. **d**, *In vitro* interaction of *Ec*ZorC with 200 nM dsDNA (52bp, 5’ FAM-labeled scrambled DNA sequence), ZorC concentrations were from lane 2 to lane 8: 1600, 100, 200, 300, 500, 1000, and 1600 nM, respectively. **e**, The effects of ZorC mutations on dsDNA binding activity, all reactions were made to a final concentration of 200 nM of dsDNA and 1000 nM of protein. Gels in **d** and **e** are representative of three independent assays. **f**, Ribbon model representation of *Ec*ZorD in complex with ATP-γ-S. *Ec*ZorD structure is shown in light purple with the bound ATP-γ-S shown in spheres. **g**, The ZorD ATP-γ-S binding site. The backbone of the DEAQ box motif (ZorD residues 730-733) is colored in magenta. Conserved negatively charged residues surrounding ZorD DEAQ box motif is shown. **h**, The effects of ZorD mutations on *Ec*ZorI-mediated anti-phage defense, as measured using EOP assays. ΔNTD represents ZorD^Δ1–502^ and ΔCTD represents ZorD^Δ503–1080^. **i**, ZorD_CTD_ degrades linear plasmid DNA. **j-l**, ZorD_CTD_ degrades phage genomic DNA (gDNA). For c and h, data represent the mean of at least 3 replicates and are normalized to the control samples lacking *Ec*ZorI. For phage Bas58, the limit of detection (LOD; one plaque observed in the assay) is marked with a bashed line. Data for additional phages are provided in **Extended Data Fig. 6a**. **6d**, **6e** and **i-l** are representative of at least 3 replicates.

### ZorC and ZorD interact with DNA

To better understand the roles of ZorC and ZorD in anti-phage defense, we next set out to obtain structural models and investigate their biological roles. ZorC is a soluble protein with no predicted function but possesses an EH signature motif (E400, H443). We determined the cryo-EM structure of *Ec*ZorC to an anisotropic resolution of 3.7 Å (**Fig. 4a, Extended Data Fig. 7a-f** and **Extended Data Table 1**). *Ec*ZorC consists of a ‘core’ domain that connects through a long linker to a C-terminal globular domain. In the core domain, we mapped the EH signature motif: E400 and H443, together with D332, R335, W339, and W458, form an electrostatic network that is buried in a pocket at the C-terminal end of the core domain (**Fig. 4b**). This network appears to be critical for Zorya function, as the individual substitution of E400 or H443 with alanine abolish Zorya defense (**Fig. 4c**). Additionally, in the AlphaFold2-predicted model, the N-terminal part of *Ec*ZorC (residues M1-E48) contains two helices, which are hydrophilic and diverge from the core domain; however, no density is observed for this region in the resolved cryo-EM map (**Extended Data Fig. 7e, g**). For the C-terminal globular domain of *Ec*ZorC, it was not possible to build an atomic model *de novo* due to its conformational flexibility. Deletion of the ZorC N-terminal two helices or of the ZorC C-terminal globular domain result in loss of Zorya function (**Fig. 4c**). Analyzing the electrostatic distribution of the hydrophilic surface of ZorC reveals several patches of positively charged regions, including the regions adjacent to the two helices and EH signature motif (**Extended Data Fig. 7h**), indicating that ZorC could interact with nucleic acids. To test this, we incubated ZorC with non-specific dsDNA oligonucleotide. We observed that ZorC migrates together with DNA, and at higher ratios of protein-DNA complex aggregation was observed, indicating that ZorC is a DNA binding protein. Deletion of the ‘cap’ domain or mutations in EH signature motif significantly alter the DNA binding activity of ZorC (**Fig. 4d, e**).

*Ec*ZorD contains 1,080 residues and has a predicted Snf2-related domain at its C-terminus^7^. This domain is known for utilizing energy derived from ATP hydrolysis to bind or remodel DNA^24^. Therefore, we determined the structure of *Ec*ZorD, in the absence and presence of ATP-γ-S (a slowly hydrolysable ATP analog), to resolutions of 2.7 Å and 2.8 Å, respectively (**Fig. 4f**, **Extended Data Fig. 8b, d** and **Extended Data Table 1**). The *Ec*ZorD N-terminal domain (residues M1-N502) interacts directly with its C-terminal domain (residues D503-A1080), forming a toroid-shaped molecule. ATP-γ-S is bound within a cleft where the hallmark DEAQ box motif (ZorD residues D730-Q733) is situated, surrounded by many conserved negatively charged residues (**Fig. 4g** and **Extended Data Fig. 8f, g**). We generated Zorya mutants that encode alanine substitutions in both the ZorD ATP binding site and those conserved negatively charged residues to assess their role in Zorya defense (**Fig. 4g**). In agreement with structural analysis, suppressing the charge of these residues resulted in loss of function of the Zorya system (**Fig. 4h**). Next, purified ZorD was incubated with plasmids to assess its DNA targeting activity. We observed that full-length ZorD is unable to degrade DNA. However, when we purified the ZorD N-terminal domain and C-terminal domain separately, we found that ZorD C-terminal domain exhibits nuclease activity that can rapidly degrade both plasmid DNA and phage genomic DNA (**Fig. 4i-4l** and **Extended Data Fig. 8a**). Mutation of the residues from the DEAQ box motif (D730A, E731A) or a glutamate (E651) recognizing the ribose sugar group of the ATP completely abolishes the nuclease activity of the ZorD C-terminal domain (**Fig.4i-4l**). These results suggest that ZorD harbors a nuclease activity, and full-length ZorD is in an autoinhibited state, which is likely activated once the defense is triggered, presumably through a conformational change.

### ZorAB recruits ZorC and ZorD during phage invasion

To examine how the ZorAB, ZorC, and ZorD activities are coordinated in response to phage infection, we explored whether it was possible to complement deletions of the *Ec*ZorI soluble components with the corresponding components of Zorya from another species. Testing a *Pseudomonas aeruginosa* type I Zorya (*Pa*ZorI) cloned under the native *Ec*ZorI promoter revealed defense activity against several of the same phages protected against by *Ec*ZorI (**Extended Data Fig. 9**). However, neither *Pa*ZorD nor *Pa*ZorCD genes could complement deletions of the corresponding *Ec*ZorI genes, hinting that direct interactions occur between at least one of ZorAB– ZorC, ZorAB–ZorD, or ZorC–ZorD (**Extended Data Fig. 9**). Our quantitative mass spectrometry data indicate there are ∼200 *Ec*ZorAB complexes (assuming most ZorA and ZorB are assembled in ZorA_5_B_2_ complexes), 155 *Ec*ZorC, and 165 *Ec*ZorD per cell possessing pZorI, implying an approximately 1:1:1 ZorA_5_B_2_:ZorC:ZorD stoichiometry (**Extended Data Fig. 2j, k** and **Extended Data Table 2**). We next used total-internal reflection fluorescence (TIRF) microscopy to examine the sub-cellular distributions of fully functional mNeonGreen (mNG) fusions to *Ec*ZorB (ZorB-mNG), *Ec*ZorC (mNG-ZorC) and *Ec*ZorD (ZorD-mNG) (**Extended Data Fig. 10a**). Expression of mNG from the *Ec*ZorI promoter resulted in uniform, cytoplasmic fluorescence independent of the presence of phage (**Extended Data Fig. 10b**). For ZorB, we observed a significant increase in the quantity of foci, that likely comprise multiple ZorAB complexes, when the bacteria were infected with Bas24 (**Fig. 5a, b**). Consistent with the anticipated integral-membrane localization, the ZorAB complexes were predominately membrane-localized and exhibited low diffusibility, irrespective of the presence of phage (**Extended Data Video 1** and **Extended Data Video 2**). In contrast, the fluorescent foci formed by the cytosolic Zorya components ZorC and ZorD freely diffused in the cytoplasm in the absence of phage (**Fig. 5c, e** and **Extended Data Video 3**). However, upon phage infection, we observed a significant increase in the number of ZorC/ZorD foci, which further appeared to become static (**Fig. 5d, f** and **Extended Data Video 4**). We presume that phage inflict damage on the cell envelope at multiple sites, which in turn activates the Zorya system to the maximum extent. The hypothesis that ZorC/D is recruited to ZorAB complexes during phage infection is reinforced by a positive correlation between the number of ZorD foci and the multiplicity of Bas24 infection (**Extended Data Fig. 10c, d**). Collectively, these results suggest that multiple ZorAB complexes detect phage infection. Subsequently, the cytosolic Zorya effectors ZorC and ZorD are recruited by ZorAB complexes to prevent phage propagation.

**Figure 5.**
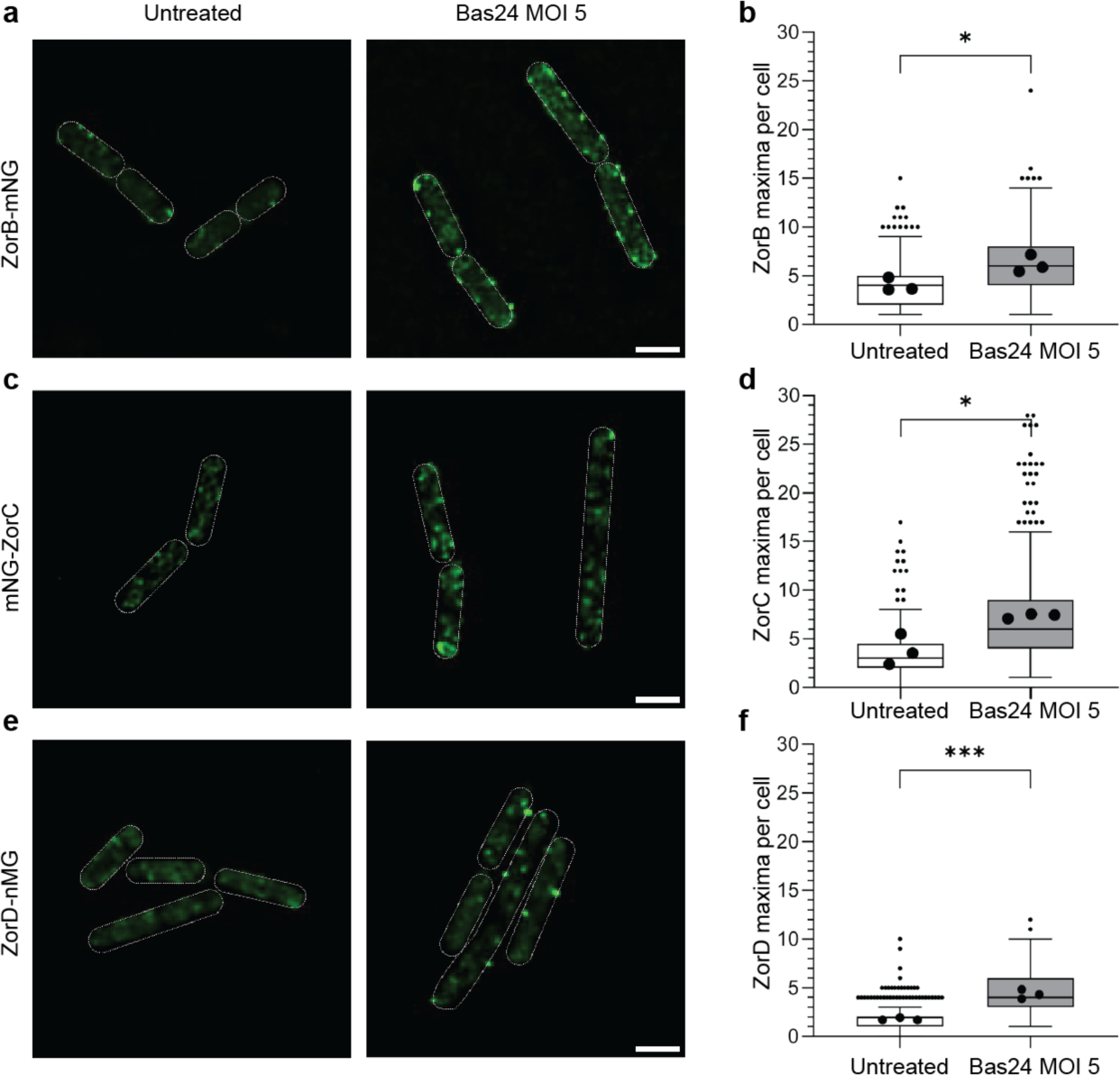
ZorAB recruits ZorC and ZorD during phage invasion. **a**, **c**, **e**, Exemplary deconvolved TIRF microscopy pictures of ZorB C-terminal, ZorC N-terminal, and ZorD C-terminal fusions with mNeongreen either untreated or exposed to Bas24 at an MOI of 5 for 30 min. **b, d, f**, Comparison of detected maxima of the ZorI proteins between untreated or exposed to Bas24 at an MOI of 5 for 30 min (n cells > 250). Means are derived from three independent biological replicates. Scale bar 2 µm. Statistics were calculated in Prism GraphPad 9 by applying the in-built analyses of unpaired t-tests or one-way ANOVA^59^. P-value: NEJM (New England Journal of Medicine) style, 0.12 ns, 0.033(*), 0.002(**), <0.001(***)

## Discussion

Here, we show that an *E. coli* type I Zorya system exhibits broad defense activity against phylogenetically diverse phages but not against bacterial conjugation or plasmid transformation. ZorA and ZorB form an inner membrane-integrated ZorA_5_B_2_ proton-driven motor complex with a long, intracellular tail structure. We propose that the ZorAB complex acts as a sensor to detect phage-induced perturbation of the cell envelope and then transmits an invasion signal, via rotation of the ZorA tail, to recruit and activate the effectors ZorC and ZorD. Activated ZorC and ZorD then bind and degrade invading phage DNA within the proximity of the infection site (**Fig. 6**).

**Figure 6.**
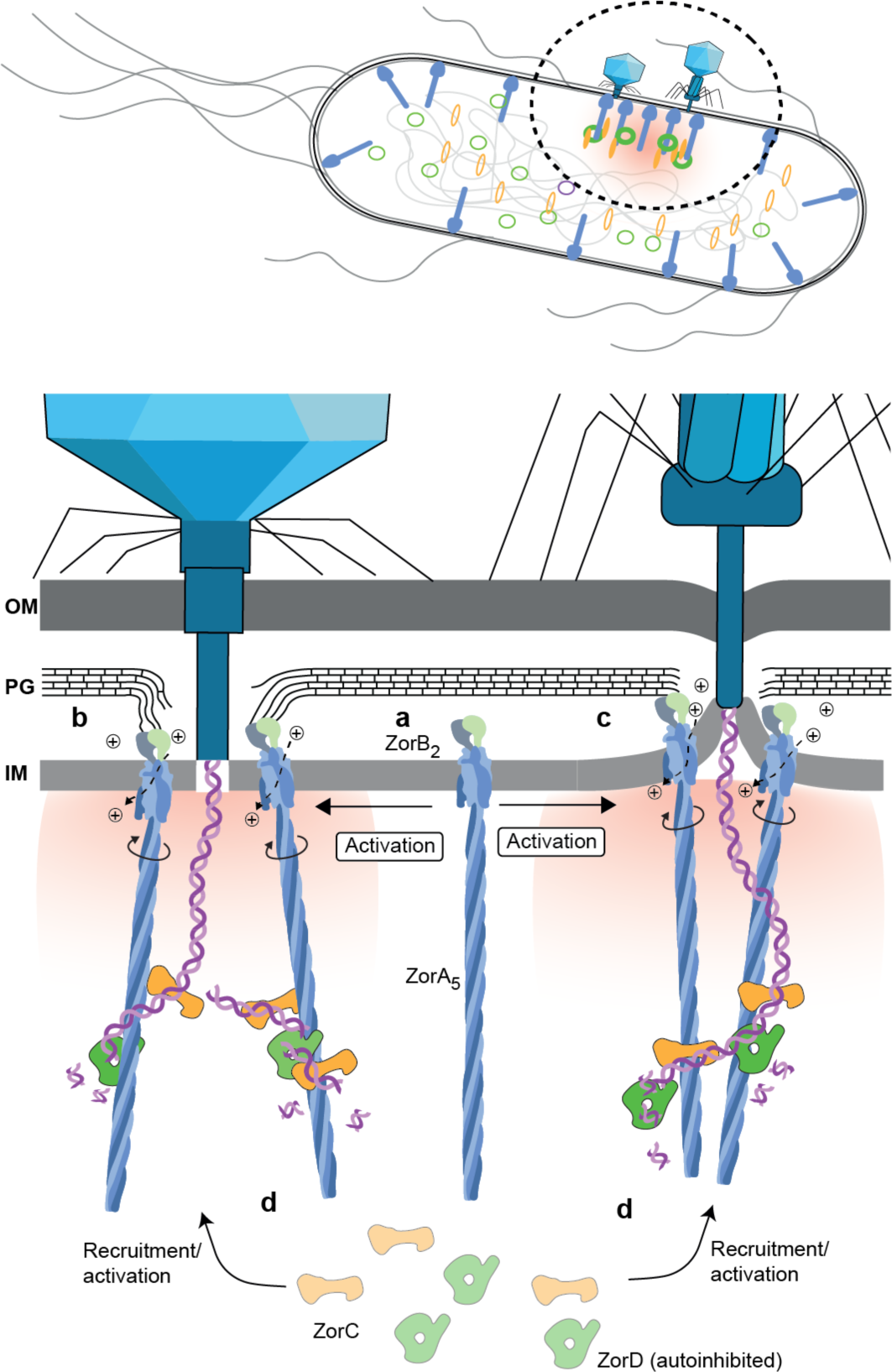
Proposed model of Zorya activation. **a**, An inactive ZorAB complex embedded in the inner membrane. **b-c**, Inactive ZorAB complex detects a signal from phage infection that (**b**) depresses the cell wall or (**c**) induces inner membrane curvature. ZorB PGBDs binds and anchors to the cell wall. Ion translocation through activated ZorAB generates rotational torque, triggering ZorA and its long intracellular tail to rotate around ZorB. **d**, The signal from the ZorAB motor is transferred through the ZorA tail, which recruits and/or activates ZorC and ZorD. Activated ZorC/D binds phage DNA, restricting it to a zone close to the site of phage genome injection, where nuclease degradation of phage DNA prevents phage infection.

Our overall model for the Zorya anti-phage defense mechanism is more likely than alternate models in which ZorC and/or ZorD function as sensors, detecting and/or digesting the phage genome and transferring this signal to the ZorAB channel. First of all, we have shown that the defense is direct and not through abortive infection, refuting a prior hypothesis of ZorAB functioning as a ZorCD–DNA activated pore. Second, ZorC and ZorD alone (without ZorAB) show no protection from phage infection. Third, neither bacterial conjugation nor plasmid transformation can activate Zorya, which would be expected if ZorC and/or ZorD act directly upon foreign DNA. Finally, flagellar stator units function in both Gram-positive and Gram-negative bacteria^15^, but Zorya is underrepresented in Gram-positive bacteria. The rare presence of Zorya in Gram-positive bacteria may be due to differences in cell wall architecture and/or phage infection mechanisms that prevent effective Zorya function, rather than a more fundamental difference prohibiting basic ZorAB rotary motor function in these bacteria.

By directly sensing perturbation of the cell envelope, Zorya would introduce an elegant mechanism that exploits the critical first stages of infection for the activation of downstream defense effectors ZorC and ZorD. Due to the distance between the inner membrane and PG layer exceeding the ‘reach’ of the ZorB PGBDs (**Fig. 3a**), ZorAB complexes would be inactive (not anchored to the PG layer) and free to diffuse laterally, as observed for MotAB complexes before flagellar incorporation^15,25^. Perturbation of the PG layer or local curvature of the inner membrane, which can occur during breaching of the cell envelope by phages^26^, would bring the ZorB PGBD closer to the PG layer, allowing binding and subsequent activation of the ZorAB motor (**Fig. 6b, c**). Our model predicts that the ‘reach’ of the ZorB PGBD should be evolutionarily diversified to adapt to different bacteria cell envelope architectures, which appears to be the case (**Extended Data Fig. 11**). Conceivably, multiple ZorAB complexes could diffuse to the sites of phage penetration, efficiently responding to the invasion signal, where they can recruit ZorCD. Confinement of ZorC and ZorD activation to the region where phage DNA injection occurs would generate a localized defense response at the sub-cellular level, potentially protecting host DNA from effector activity (ZorC/D) without the need of epigenetic-based self vs. non self-discrimination mechanisms^27^. Such local defense also saves energy, as it is only activated when needed, and could allow the Zorya system to rapidly turn off again after effective defense.

In addition to the mechanistic insights directly supported by our data, there are several potential aspects worthy of future investigations. A first open question is the role of two striking features of the ZorAB structure: the rotary motor and the obligatory long tail. One possible explanation for the role of rotation is in signal transduction from phage ingress at the cell envelope to activation of ZorC and ZorD. If the direction of the ZorAB motor is the same as for MotAB (clockwise rotation of A around B, as viewed from the outside of the cell), rotation could potentially induce untwisting of the pentameric ZorA tail super-helix, thereby altering the conformation of the tail to facilitate recruitment or activation of ZorC and/or ZorD (**Extended Data Fig. 12a, b**). A somewhat similar sensory transmission mechanism occurs with bacterial chemosensory arrays and the long ZorA tail might serve to transmit the signal in a manner analogous to the long coiled-coil cytoplasmic signaling domains of methyl-accepting chemotaxis proteins^23^. A second hypothesis that would explain the need for both rotation and a long ZorA tail is that rotation might allow the ‘reeling in’ of the phage DNA around the tail (**Extended Data Fig. 12c**). This would constrain phage DNA to the entry site and keep it in an inactive state (e.g. preventing RNA polymerase access), resolving the time pressure for ZorC and ZorD recruitment and activation, and nuclease activity to take effect before the host takeover mechanisms (see **Extended Data Fig. 12** and **Legend Discussion**). This would also change the required timescale of the defense, giving the host a fighting chance against phages which often have extremely fast lifecycles. Finally, a long, rotating, flexible tail could increase the possibility of catching phage DNA for degradation.

Another interesting open question is that we observed in our microscopy experiments the increase in Zorya-mNG foci and intensity for cells infected with Bas24, against which *Ec*ZorI shows protection, as well as T4 (**Extended Data Fig. 10e**), against which it does not. This indicates that the system is activated by both phages, in agreement with the hypothesis that the piercing of the cell envelope is the triggering signal. However, we do not yet know why, despite ZorAB activation, phage T4 is not susceptible to *Ec*ZorI defense (**Fig. 1b**), suggesting T4 resists ZorC or ZorD activity, potentially via a T4-encoded anti-Zorya protein. Phage-encoded proteins that inactivate Zorya effectors may be one reason for the existence of multiple Zorya types, each encoding the ZorAB core but with different effectors^7^.

In summary, we provide structural and functional insight into the Zorya defense system and propose that Zorya functions as a direct defense system against a wide range of phages, which acts during the early infection process by initiating a localized response within the proximity of the phage injection site. Our studies reveal a unique activation signal of an anti-phage defense system and pave the way for further studies to understand its detailed mechanism.

## Methods

### Phylogenetic analysis of Zorya systems

To create the phylogenetic tree shown in **Fig. 1a**, the operons that encode the Zorya (Zor) system types I, II, and III were download from the PADLOC server^8,28^. As ZorA and ZorB are present in all Zorya types, the tree was generated using concatenated ZorA+B sequences. MMseqs2^29^ was used to remove the redundancy with the parameters ‘–min-seq-id 0.90’ and ‘-c 0.8’. The filtered sequences were then aligned using MAFFT^30^ with parameters: --maxiterate 1000 –globalpair. The alignment was then trimmed using trimAl^31^ with the parameter: -gt 0.25. The phylogenetic tree was constructed using IQ-TREE^32^ with the parameters -nstop 500 -bb 2000 -m LG+G4. iTOL^33^ was used for tree display and annotation.

### Cloning of Zorya defense system and mutagenesis

The DNA of *E. coli* ZorI full operon was amplified from the *E. coli* strain NCTC9026 genome (purchased from the National Collection of Type Cultures, NCTC) with its native promotor and was subcloned into a modified pACYC vector using In-Fusion cloning strategy (In-Fusion® Snap Assembly Master Mix; TaKaRa Cat. # 638947). The DNA of *P. aeruginosa* ZorI full operon was amplified from the *P. aeruginosa* strain DSM24068 genome (DSMZ-German Collection of Microorganisms and Cell Cultures GmbH; Leibniz Institute) and was subcloned into a modified pACYC vector under *E. coli* ZorI native promotor using In-Fusion cloning strategy. For generating mutations (point mutations, deletions, mNeonGreen tag insertion, *Ec*ZorI ZorC or ZorCD genes replaced by *Pa*ZorI ZorC or ZorCD genes), plasmids were constructed based on standard cloning techniques (In-fusion snap assembly (TakaraBio)). All plasmids were verified by either Sanger or nanopore sequencing.

### Phage infectivity assays

The host *E. coli* MG1655 ΔRM^19^ possessing either pControl (pACYC) or p*Ec*ZorI (or mutants thereof) were grown overnight in LB + Chloramphenicol (Cm; 25 µg/mL). Efficiency of plaquing (EOP) assays were performed using bacterial lawns of the host strain in 0.35% LB agar + 10 mM MgSO_4_ + 2 mM CaCl_2_ overlaid onto 1.5% LB agar + Cm. Ten-fold dilution series of phages were spotted onto the overlays, air-dried, then the plates were incubated overnight at 30°C. Liquid culture infection time courses were performed in 96-well plates in an incubated shaking plate reader at 30°C. The time courses were begun with cells at an OD_600_ of 0.05 and phages were added at the indicated MOI, assuming an OD_600_ to cell ratio of 3×10^8^ cells per OD_600_ unit.

### Phage adsorption and one-step growth curves assays

Overnight cultures of *E. coli* MG1655 ΔRM possessing either pControl or p*Ec*ZorI were used to inoculate fresh LB + Cm cultures at a 1:100 dilution. The inoculated cultures were grown at 30 °C with shaking until reaching an OD_600_ of 0.4–0.6, then harvested by centrifugation, washed with LB + Cm, and resuspended at an OD_600_ of 1.0 in LB + 10 mM MgSO_4_ + 2 mM CaCl_2_. For phage the adsorption assays, 10 mL samples of resuspended cells were infected with phage Bas24 at an MOI of 10^−4^, then the samples were mixed and incubated at 30°C without shaking. For the 0 min timepoint (total input phages), 100 µL samples were removed and added to 0.35% LB Agar seeded with *E. coli* MG1655 ΔRM + pControl (as an indicator lawn), then poured on top of 1.5% LB agar + Cm. For each subsequent time point, 1 mL samples were taken, centrifuged to pellet cells, then the supernatant (containing unabsorbed phages) was filtered through a 0.2 µm PES syringe filter. Samples (100 µL) of the filtered supernatant were added to indicator overlays (as above) poured onto 1.5% LB agar + Cm. All overlay plates were incubated overnight at 30°C before counting plaques. For each timepoint, the percentage of unabsorbed phages was calculated as the timepoint plaque count / plaque count for the time 0 min pControl sample. For the one-step phage growth curves (burst time and size), 2 mL samples of the cells resuspended at and OD_600_ of 1.0 in LB + 10 mM MgSO_4_ + 2 mM CaCl_2_ (as above) were infected with phage Bas24 at an MOI of MOI of 10^−4^, then the samples were mixed and two 10-fold diluted samples were prepared, then the dilution series for each sample was incubated at 30°C without shaking. At the indicated timepoints, 100 µL samples of each dilution were removed and added to 0.35% LB Agar seeded with E. coli MG1655 ΔRM + pControl (as an indicator lawn), then poured on top of 1.5% LB agar + Cm. All overlay plates were incubated overnight at 30°C before counting plaques. For each timepoint, the plaque forming units (PFU) were normalized to the PFU of the 0 min pControl samples.

### Conjugation assays

Plasmids encoding kanamycin (Km) resistance and each possessing different origins of replications (ColE1;pMAT16, RSF1010;pPF1825^34^, pBBR1;pSEVA237R^35^, or RK2;pPF1619) were conjugated from the *E. coli* donor ST18^36,37^(auxotroph requiring supplementation with 5-aminolevulinic acid; ALA) into the *E. coli* recipient MG1655 ΔRM possessing either pControl or p*Ec*ZorI. Mattings were performed at the indicated donor to recipient ratios and incubated overnight on LB agar + Cm + ALA at 30 °C. The conjugation efficiency was determined by plating dilution series of the matings onto LB agar + Cm + Km (transconjugants) and LB agar + Cm (total recipients). The transconjugant frequency was defined as the transconjugant CFU/recipient CFU.

### Transformation assays

Chemically competent cells of *E. coli* MG1655 ΔRM possessing either pControl or p*Ec*ZorI were prepared by the Inoue method^38^, with HEPES-KOH pH 6.8 used for the transformation buffer. Cells were stored in 200 µL aliquots at −80 °C prior to use. For each transformation assay, 5 ng of plasmid (quantitated using a Qubit BR kit) was used. Plasmids used were as above for the conjugation assays (ColE1;pMAT16, or pBBR1;pSEVA237R).

### Cell survival assays

Overnight cultures of *E. coli* MG1655 ΔRM possessing either pControl or p*Ec*ZorI were used to inoculate fresh LB + Cm cultures at a 1:100 dilution. The inoculated cultures were grown at 30 °C with shaking until reaching an OD_600_ of 0.4–0.6, then harvested by centrifugation, washed with LB + Cm, and resuspended at an OD_600_ of 0.2. Phage Bas24 was then added at an MOI of 5 to each sample; control samples without phage addition were also included. After 20 min adsorption, 10-fold serial dilutions of each sample were plated (100 µL each) on LB + Cm, then incubated overnight at 30°C. The cell survival rate was calculated as the CFU obtained + Bas24/CFU obtained without phage addition.

### Phages and phage genome purification

Phage primary stocks were prepared using the double-agar method^39^ by setting the grow conditions to almost confluent plaques. The phages were collected by adding SM buffer (100 mM NaCl, 8 mM MgSO_4_, 50 mM Tris-HCl, pH 7.5, 5mM CaCl_2_) on top of the overly agar and mixed for 4 h at 4 °C. The suspension was collected and centrifuged 15 min at 4000 g. High titer phage samples were obtained by inoculating 1-3 L of LB with a 10^3^ dilution of an overnight culture of MG1655 ΔRM and grown at 37 °C to and OD_600_ of 0.3. The bacterial culture was inoculated with the primary stock to an MOI of 0.025 and infection was carried out at 37 °C at 90 rpm until a clear lysate was obtained. The lysate was harvested at 4000 g, 15 min and 4 °C. After decanting the supernatant, 1 µg/mL of DNase I and 1 µg/mL of boiled RNase A were added to the cleared lysate. The lysate was gently stirred at 90 rpm for 30 min at room temperature (RT).

Phages were concentrated by polyethylene glycol (PEG) precipitation. NaCl was gradually added to a final concentration of 1 M, followed by gradual addition of 10% PEG 8,000 with continuous stirring at RT until dissolved. After obtaining a clear solution, the lysate was stirred (100 rpm, 30 min; 4°C) and left overnight at 4°C. The lysate was centrifuged (15,000 g, 1 h, 4°C) and the clear supernatant was removed. The precipitate was resuspended in the minimal amount (up to 2 ml) of SM buffer that allowed solubilization. Insoluble materials were removed by adding 20% v/v of chloroform and centrifuged (8,000 g, 10 min). The supernatant was stored at 4°C to be used as phage sample for the following step. The phage was then purified by rate zonal separation using OptiPrepTM Density Gradient Medium (Sigma Aldrich) density gradient ranging from 50 to 10%, diluted in SM media. Phage sample was applied on the top of the gradient and centrifuged (150,000 g, 18 h, 4°C). The phage was extracted, dialyzed against SM buffer and samples were stored at 4°C. The phage genomes were extracted using the Phage DNA isolation kit from Norgen Biotek, aliquoted and stored at −20°C.

### Protein expression and purification

#### ZorAB

The full-length genes of *E. coli* ZorA and ZorB code for 729 and 246 residues, respectively. The tandem gene was PCR amplified from the *E. coli* strain NCTC9026 genome (purchased from the National Collection of Type Cultures, NCTC) and subcloned into a modified pET vector containing a C-terminal human rhinovirus (HRV) 3C protease cleavage site and a twin-Strep-tag II (resulting in pET11a-ZorA-ZorB-3C-TSII). The plasmids containing the recombinant genes were transfected into *E. coli* C43(DE3) competent cells and the proteins were expressed in LB medium. When the culture OD reached to 0.6-0.8, the temperature was decreased from 37°C to 24°C (OD reached approximately 0.8-1.0), and 0.5 mM isopropyl β-D-1-thiogalactopyranoside (IPTG) was added for overnight protein induction. The culture was harvested, and the cell pellet was resuspended in buffer A containing 20 mM HEPES-NaOH pH 7.5, 300 mM NaCl supplemented with EDTA-free protease inhibitor (Thermo Fisher Scientific) and lysozyme from chicken white egg (Sigma) to a final concentration of 50 μg/mL and Deoxyribonuclease I from bovine (Sigma) to a final concentration of 30 μg/mL. The mixture was disrupted by high-pressure homogenizer and spun at 185,000 g for 1 h. The pellet containing the membrane was collected and was solubilized using buffer B containing 30 mM HEPES-NaOH pH7.5, 300 mM NaCl, 10% glycerol, 2% Lauryl Maltose Neopentyl Glycol (LMNG; Anatrace), supplemented with EDTA-free protease inhibitor at 4°C for 2 h. The solubilized membrane was then spun at 90,000 g rpm for 40 min and the supernatant was loaded onto a gravity flow column containing 2 mL (resin volume) of Strep-Tactin® Superflow® high-capacity resin (IBA), pre-equilibrated with wash buffer containing 20 mM HEPES-NaOH pH 7.5, 300 mM NaCl, 10% glycerol and 0.005% LMNG. The resins were washed five times with 2-3 resin volumes of the wash buffer and elution was carried out five times with 0.5 resin volume (1 mL) of elution buffer containing 20 mM HEPES-NaOH pH 7.5, 300 mM NaCl, 10% glycerol, 0.005% LMNG and 10 mM desthiobiotin). The recombinant protein was then concentrated and loaded onto a pre-equilibrated (20 mM HEPES-NaOH pH 7.5, 150 mM NaCl, 0.002% LMNG) Superose 6 Increase 10/300 GL Size-exclusion chromatography column. Fractions from the elution peak corresponding to the molecular weight of ZorAB complex were pooled, and the protein was concentrated for cryo-EM grids preparation and functional experiments. The procedures of expression and purification of ZorAB mutants were similar as the ZorAB wild type.

#### ZorC

The predicted ZorC gene codes for 560 residues. The ZorC gene together with a short region upstream of ZorC N-terminus that codes for 7 residues (LPVGYAT) was PCR amplified from the DNA genome of *E. coli* strain NCTC9026 and subcloned into the modified pET vector (resulting in pET11a-ZorC-3C-TSII). *E. coli* BL21 (DE3) gold chemically competent cells were transformed with the plasmids and the protein was expressed in LB medium with the presence of 100 μg/mL of ampicillin. Briefly, when the OD reached to 1.0-1.2, decreased the temperature to 16°C and 0.5 mM isopropyl β-D-1-thiogalactopyranoside (IPTG) was added for overnight protein induction. The culture was harvested, and the cell pellet was resuspended using buffer containing 20 mM Tris pH 7.5, 10% glycerol and 500 mM NaCl supplemented with EDTA-free protease inhibitor (Thermo Fisher Scientific). The cells were lysed using an Avestin Emulsiflex C3 homogeniser, cooled to 4°C, and spun at 18,000 g for 40 minutes. The supernatant was then added to a gravity flow column containing 3 mL (resin volume) of Strep-Tactin® Superflow® high-capacity resins (IBA), pre-equilibrated with wash buffer (20 mM Tris pH 7.5, 10% glycerol and 500 mM NaCl). Resins were washed five times with 2-3 resin volumes of wash buffer and elution was performed with 4 CV of elution buffer (20 mM Tris-HCl pH 7.5, 500 mM NaCl and 10 mM desthiobiotin). The recombinant protein was pooled and concentrated and was loaded onto a pre-equilibrated (20 mM Tris-HCl pH 7.5, 500 mM NaCl) Superose 6 Increase 10/300 GL Size exclusion chromatography column. Peak fractions were pooled, and another round of size exclusion chromatography was carried out with buffer 20 mM HEPES-NaOH pH 7.5, and 150 mM NaCl to decrease NaCl concentration. Fractions from the elution peak corresponding to the molecular weight of ZorC were pooled and the protein was concentrated to approximately 1 mg/mL for cryo-EM grids preparation and functional experiments. ZorC proteins used for electromobility shift assays (EMSAs) was kept in elution buffer B (20 mM Tris-HCL pH 7.5, 500 mM NaCl, 10 % glycerol) and flash frozen in small aliquots and stored at −80C until use. Sample purity was assessed by SDS-PAGE. The procedures of expression and purification of ZorC mutants were similar to those for the ZorC wild type.

#### ZorD

The predicted ZorD gene coding 1,086 residues was PCR amplified from the DNA genome of *E. coli* strain NCTC9026 and was subcloned into the modified pET vector. The expression and purification of ZorD protein were similar to ZorC protein, except for the purification buffer. Briefly, the suspension buffer contained 150 mM NaCl and 20 mM HEPES at pH 7.5, 10 glycerol; the wash buffer was the same as the suspension buffer and the elution buffer contained 150 mM NaCl and 20 mM HEPES-NaOH at pH 7.5, 10% glycerol with 10 mM desthiobiotin; and the size exclusion chromatographic buffer contained 20 mM HEPES-NaOH pH 7.5 and 150 mM NaCl. Purified ZorD was concentrated to 0.4-0.6 mg/mL for functional experiments and cryo-EM grids preparation. For nuclease experiment, ZorD protein was kept in the elution buffer and flash frozen in small aliquots and stored at −80**°**C until use.

### Cryo-EM grid preparation, data collection, model building, and refinement

#### ZorAB

Freshly purified ZorAB sample was concentrated to 2-3 mg/mL and 2.7 μL protein was applied onto the glow-discharged (30 s, 5 mA) grids (Quantifoil R0.6/1 300 mesh Au) and plunge-frozen into liquid ethane using a Vitrobot Mark IV (FEI, Thermo Fisher Scientific), with the settings: 100% humidity, 4°C, blotting force 25, 4-6 s blot time and 7 s wait time; Movies were collected using the semi-automated acquisition program EPU (FEI, Thermo Fisher Scientific) on a Titan Krios G2 microscope operated at 300 keV paired with a Falcon 3EC direct electron detector (FEI, Thermo Fisher Scientific). Images were recorded in electron counting mode, at 96,000x magnification with a calibrated pixel size of 0.832 Å and an underfocus range of 0.7 to 2.5 μm. The number of micrographs and total exposure values for the different datasets are summarized in Table S1. Grids preparation, and data collection strategies of the ZorAB mutants were similar to those for the ZorAB wild type.

#### ZorC

Purified ZorC (3 μL at ∼1 mg/mL) was applied onto glow-discharged (30 s, 5 mA) grids (UltrAuFoil R 0.6/1, 300 mesh, Gold) and plunge-frozen into liquid ethane using a Vitrobot Mark IV (FEI, Thermo Fisher Scientific), with the settings: 100% humidity, 4 °C, blotting force 20, 4 s blot time and 10 s wait time; Movies were collected using the semi-automated acquisition program EPU (FEI, Thermo Fisher Scientific) on a Titan Krios G2 microscope operated at 300 keV paired with a Falcon 3EC direct electron detector (FEI, Thermo Fisher Scientific). Images were recorded in electron counting mode, at 96,000x magnification with a calibrated pixel size of 0.832 Å and an underfocus range of 1 to 2.5 μm. The number of micrographs and total exposure values for the datasets are summarized in **Extended Data Table 1**.

#### ZorD

ZorD showed preferred orientation of particles on ice. 0.5% zwitterionic detergent CHAPSO (Anatrace) was added to the purified sample to a final concentration of 0.0125% before cryo-EM grid preparation. For the apo form, the preparation of grids was similar to ZorC. For ZorD in complex with ATP-γ-S, 4 µL of 0.1 mM ATP-γ-S was added into 400 µL of purified ZorD at 0.0375 mg/ml. The mixture was concentrated to 15 µL to reach a ZorD concentration of around 0.6 mg/mL. The grid preparation was similar to ZorC. The number of micrographs and total exposure values for the different datasets are summarized in **Extended Data Table 1**.

### Cryo-EM data processing

All datasets were processed using cryoSPARC^40^ v4.2.1, unless otherwise stated. We started by using Patch motion correction to estimate and correct for full-frame motion and sample deformation (local motion). Patch Contrast function (CTF) estimation was used to fit local CTF to micrographs. Micrographs were manually curated to remove the relatively ice thickness value bigger than 1.1 and the CTF value worse than 3.5 Å. We performed particle picking by template picking or using topaz particle picking^41^. Particles were extracted with a box size of 500 pixels for ZorAB datasets, 256 pixels for the ZorC dataset, and 400 pixels for the ZorD dataset. One round of 2D classification was performed followed by *ab initio* reconstruction. Heterogeneous refinement was used to exclude broken particles. Non-uniform refinement was applied with a dynamic mask to obtain a high-resolution map. Local refinement was additionally performed with a soft mask to achieve a higher-resolution map of some flexible regions. For all datasets, the number of movies, the number of particles used for the final refinement, map resolution, and other values during data processing are summarized in the **Extended Data Table 1**.

### Model building and validation

We used AlphaFold2^42^ (AF2) to predict all the initial models. The predicted models were manually fit into the cryo-EM density by using UCSF ChimeraX^43^. The model was refined in Coot^44^ or refined using StarMap^45^ in the case of ZorC, for which the map is anisotropic and the resolution is modest. The model was then refined against the map using PHENIX real space refinement^46^. The ZorAB composite model was constructed by extending the pentameric tail as an idealized right-handed super-helical coiled coil. Local conformations were manually adjusted in PyMol^47^(v2.5) and optimized through energy minimization using Gromacs (v2022.5)^48^. However, it’s worth noting an irregularity in the AF2 model, specifically in residues 312 to 322, which introduces a significant twist in the tail, raising possibilities of other pentameric forms of the ZorA tail and further reflecting its conformational dynamics.

### Peptidoglycan (PG) purification and pull-down experiment

Peptidoglycan was purified from *E. coli* strain MG1665 ΔRM with the protocol adapted from^49^ Briefly, *E. coli* strain MG1665 ΔRM cells were incubated in 1 L LB media until the OD_600 nm_ reached 0.8. Cells were harvested and resuspended in 12 mL PBS buffer and split into two 50 mL falcon tubes, SDS solution was added to final the concentration of 6%. The falcon tubes were boiled for 1 hour while stirred at 500 rpm. The heat was turned off and the tube was allowed to cool to ambient temperature overnight. The next day, the solutions from both Falcon tubes were pooled into one 50 mL Falcon tube, and centrifuged at room temperature for 45 mins at 108,000 g. The pellet was washed five times with 5 mL Milli-Q® water. The PG was resuspended in 20 mL of buffer containing 50 mM Tris-HCl, pH 7.0, and α-amylase was added (SigmaAldrich) to a final concentration of 100 μg/mL and incubated for 2 hours at 37°C. Next, 50 μg/mL RNase A (Roche) and 10 μg/mL DNase (Sigma-Aldrich) were added and incubated for an additional 2 hours at 37°C. Then, the mixture was supplemented with 20 mM MgSO_4_, 10 mM CaCl_2_, and 100 μg/mL trypsin (SigmaAldrich), and incubated at 37°C overnight. The following day, EDTA at pH 8 was added to a final concentration of 10 mM to the mixture and SDS solution to a final concentration of 1%. The mixture was boiled for 20 mins in a water bath and allowed to cool to ambient temperature. The tube was centrifuged at 108,000 g for 45 minutes. The resulting pellet was washed five times with Milli-Q® water to remove residual SDS. Finally, the pellet was resuspended in 300 μL of Milli-Q® water, aliquoted into 35 μL portions, and stored at −20 °C. For PG pull-down assays, the PG was washed with 1 mL PBS+0.002% LMNG buffer and centrifuged at 20,000 g for 30 minutes. 10 μL of purified ZorAB was incubated (at a concentration of 2 mg/mL) with the PG at room temperature for 1 hour, then centrifuged at 20,000 g for 30 minutes, the supernatant was retained for SDS gel analysis. The pellet was resuspended with 20 μL of buffer and 5 μL of loading dye was added for SDS gel analysis.

### ZorC DNA-binding experiments

#### Electromobility shift assays

Frozen aliquots of ZorC and ZorC mutants in Elution buffer B (20 mM Tris-HCL pH 7.5, 500 mM NaCl, 10% glycerol) were thawed, and centrifuged to remove aggregates and diluted to 2 μM in phosphate-buffer saline (PBS) (15.2 mM NaHPO_4_, 0.90 mM calcium chloride, 2.7 mM potassium chloride, 1.47 mM potassium dihydrogen phosphate, 8.1 mM sodium hydrogen phosphate, 0.49 mM magnesium chloride, 137.9 mM sodium chloride at pH 6.8). Scrambled dsDNA substrates (5’-FAM-CTAGAAAGACTTTTAACAGTGGCCTTATTAAATGACTTCTCAACCATCTTGCTGA-3’) were incubated with a protein in PBS. All components were incubated at 4°C for 10 min. Glycerol was added to the reactions prior to loading on a 2% DNA agarose gel. Samples were run for 15-25 minutes at 250 V, and the gels were visualized using Odyssey^®^ XF Imaging System at wavelength 600 nm.

### Nuclease assays

ZorD was incubated with 200 ng pUC19 (linearized by KpnI (NEB) enzyme) in the reaction buffer containing 1 × Cutsmart buffer (NEB), 2 mM ATP (NEB) in a total volume of 25 μl. Mixture was incubated at 37°C for 1 hour and shaken at 600 rpm using an Eppendorf ThermoMixer. DNA product was purified using a NucleoSpin Gel and PCR Clean-up kit (Machery Nagel) using the standard protocol and was analyzed with 1% E-Gel™ EX. For the reaction with the phage genomes, 200 nM Proteins were incubated with around 100 ng purified phage genomic DNA in the same reaction buffer indicated above. The reactions were terminated by adding 1× E-gel loading buffer and product was analyzed with 1% E-Gel™ EX.

### Mass spectrometry sample preparation

Overnight cultures *of E. coli* MG1665 ΔRM, transformed with plasmids encoding the Zorya operon (or control), were used to inoculate (at a 1:1000 dilution) 3 mL LB media with antibiotics, then grown to an OD_600_ of approximately 0.4. The cell pellet was collected, resuspended in 500 μL of 0.2 M Tris-HCl pH 8.0, and incubated for 20 min. 250 μL of buffer containing 0.2M Tris-HCl pH 8.0, 1 M sucrose, and 1 mM EDTA, was added into the solution along with 3 μL of 10 mg/mL lysozyme. The mixture was incubated for 30 minutes, and 250 μL of 6% SDS was added to a final concentration of 1%, after which the sample was heated to 99°C for 10 min. The mixture was sonicated to fragment DNA and RNA.

For MS analysis, we performed Protein Aggregate Capture digestion of proteins^50^. To this end, 250 μL of bacterial lysate was taken from the total sample, and 750 μL of acetonitrile was added into the mixture, along with 50 µL magnetic microspheres that had been prewashed with PBS buffer. The mixture was allowed to settle for 10 min, prior to retention of the magnetic microspheres by magnetic plate. Beads were washed once with 1 mL acetonitrile, and once with 1 mL of 70% ethanol, after which all ethanol was removed and beads were stored at −20°C until further processing. Frozen beads were thawed on ice, supplemented with 100 µL ice-cold 50 mM Tris-HCl pH 8.5 buffer supplemented with 2.5 ng/µL Lys-C, and gently mixed (on ice) every 5 min for 30 min. Digestion was performed for 3 h using a Eppendorf ThermoMixer shaking at 1,250 rpm at 37 °C. Following this, beads were chilled on ice, and 250 ng of sequencing-grade trypsin was added, after which samples were gently mixed (on ice) every 5 min for 30 min. Final digestion was performed overnight using a Eppendorf ThermoMixer shaking at 1,250 rpm at 37°C. Peptides were separated from magnetic microspheres using 0.45 µm filter spin columns, and peptides were reduced and alkylated by adding TCEP and chloroacetamide to 5 mM for 30 min prior to peptide clean-up via low-pH C18 StageTip procedure. C18 StageTips were prepared in-house, by layering four plugs of C18 material (Sigma-Aldrich, Empore SPE Disks, C18, 47 mm) per StageTip. Activation of StageTips was performed with 100 μL 100% methanol, followed by equilibration using 100 μL 80% acetonitrile (ACN) in 0.1% formic acid, and two washes with 100 μL 0.1% formic acid. Samples were acidified to pH <3 by addition of trifluoroacetic acid to a concentration of 1%, after which they were loaded on StageTips. Subsequently, StageTips were washed twice using 100 μL 0.1% formic acid, after which peptides were eluted using 80 µL 30% ACN in 0.1% formic acid. All fractions were dried to completion using a SpeedVac at 60 °C. Dried peptides were dissolved in 25 μL 0.1% formic acid (FA) and stored at −20 °C until analysis using mass spectrometry (MS).

Approximately 1 µg of peptide was analyzed per injection. All samples were analyzed on an EASY-nLC 1200 system (Thermo Fisher Scientific) coupled to an Orbitrap™ Astral™ mass spectrometer (Thermo Fisher Scientific). Samples were analyzed on 20 cm long analytical columns, with an internal diameter of 75 μm, and packed in-house using ReproSil-Pur 120 C18-AQ 1.9 µm beads (Dr. Maisch). The analytical column was heated to 40 °C, and elution of peptides from the column was achieved by application of gradients with stationary phase Buffer A (0.1% FA) and increasing amounts of mobile phase Buffer B (80% ACN in 0.1% FA). The primary analytical gradient ranged from 10 %B to 38 %B over 57.5 min, followed by a further increase to 48 %B over 5 min to elute any remaining peptides, and by a washing block of 15 min. Ionization was achieved using a NanoSpray Flex NG ion source (Thermo Fisher Scientific), with spray voltage set at 2 kV, ion transfer tube temperature to 275 °C, and RF funnel level to 50%. All full precursor (MS1) scans were acquired using the Orbitrap™ mass analyzer, while all tandem fragment (MS2) scans acquired in parallel using the Astral™ mass analyzer. Full scan range was set to 300-1,300 m/z, MS1 resolution to 120,000, MS1 AGC target to “250” (2,500,000 charges), and MS1 maximum injection time to “150”. Precursors were analyzed in data-dependent acquisition (DDA) mode, with charges 2-6 selected for fragmentation using an isolation width of 1.3 m/z and fragmented using higher-energy collision disassociation (HCD) with normalized collision energy of 25. Monoisotopic Precursor Selection (MIPS) was enabled in “Peptide” mode. Repeated sequencing of precursors was minimized by setting expected peak width to 20 s, and dynamic exclusion duration to 20 s, with an exclusion mass tolerance of 10 ppm and exclusion of isotopes. MS2 scans were acquired using the Astral mass analyzer. MS2 fragment scan range was set to 100-1,500 m/z, MS2 AGC target to “50” (5,000 charges), MS2 intensity threshold to 50,000 charges per second, and MS2 maximum injection time to 5 ms; thus requiring a minimum of 250 charges for attempted isolation and identification of each precursor. Duty cycle was fixed at 0.3 s, acquiring full MS scans at ∼3.3 Hz and with auto-fitting of Astral scans resulting in MS2 acquisition at a rate of ∼100-200 Hz.

### Mass spectrometry data analysis

All RAW files were analyzed using MaxQuant software (v2.4.3.0)^51^, the earliest release version to support Astral RAW files. Default MaxQuant settings were used, with exceptions outlined below. For generation of the in silico spectral library, the four full-length Zorya protein sequences were entered into a FASTA database, along with all (23,259) Swiss-Prot-reviewed *E. coli* sequences (taxonomy identifier 562) downloaded from UniProt^52^ on the 7th of September, 2023. The data was first searched using pFind (v3.2.0)^53^, using the “Open Search” feature to determine overall peptide properties and commonly occurring (affecting >1% of PSMs) peptide modification in an unbiased manner. For searching Astral .RAW files using pFind, .RAW files were first converted to .mzML using OpenMS (v3.0.0)^54^. For the main data search using MaxQuant, digestion was performed using “Trypsin/P” with up to 2 missed cleavages (default), with a minimum peptide length of 6 and a maximum peptide mass of 5,000 Da. No variable modifications were considered for the first MS/MS search, which is only used for precursor mass recalibration. For the MS/MS main search a maximum allowance of 3 variable modifications per peptide was set, including protein N-terminal acetylation (default), oxidation of methionine (default), deamidation of asparagine, peptide N-terminal glutamine to pyroglutamate, and replacement of three protons by iron (cation Fe[III]) on aspartate and glutamate. Unmodified and modified peptides were stringently filtered by setting a minimum score of 10 and 20, and a minimum delta score of 20 and 40, respectively. First search mass tolerance was set to 10 ppm, and maximum charge state of considered precursors to 6. Label-free quantification (LFQ) was enabled, “Fast LFQ” was disabled. iBAQ was enabled. Matching between runs was enabled with a match time window of 1 min and an alignment time window of 20 min. Matching was only allowed between same-condition replicates. Data was filtered by posterior error probability to achieve a false discovery rate of <1% (default), at the peptide-spectrum match, protein assignment, and site-decoy levels.

### Mass spectrometry data statistics

All statistical data handling was performed using the Perseus software^55^, including data filtering, log2-transformation, imputation of missing values (down shift 1.8 and width 0.15), and two-tailed two-sample Student’s t-testing with permutation-based false discovery rate control. In order to determine relative concentration of all proteins in the samples, LFQ-normalized intensity values for each protein were adjusted by molecular weight. To approximate absolute copy-numbers, we extracted known protein copy-numbers based on the “LB” condition as reported by Schmidt et al.,^56^ log2-transformed them, and aligned them to the molecular-weight adjusted LFQ intensity values from our own data, resulting in 1,901 out of 2,418 quantified protein-groups receiving a known copy-number value (R² = 0.6129). Next, we subtracted the overall median from all log2 values, and determined the absolute delta between the values of each pair. Out of all pairs, 459 had a log2 delta of <0.5, which we considered as a “proteomic ruler”. Linear regression was performed on the remaining pairs (R² = 0.9868) to determine a conversion factor between MW-adjusted LFQ intensity and absolute copy-number.

### TIRF microscopy cultivation conditions

Overnight cultures of *E. coli* strains expressing ZorB, C or D translational fusions to mNeongreen were incubated shaking at 180 rpm in LB Lennox containing 20 mM MgSO_4_, 5 mM CaCl_2_ and supplemented with 12.5 µg/ml chloramphenicol at 30 °C. On the next day, a sub-culture was inoculated 1:100 and grown at 30 °C until an OD600 between 0.3-0.5 was reached. Subsequently, cells were diluted to an OD600 of 0.2 to ensure reproducible ratios between bacteria and phages. Cells were then exposed to phages at indicated MOIs or incubated untreated for 30 min in a 2 mL Eppendorf tube under shaking conditions (<650 rpm in an Eppendorf ThermoMixer). For TIRF microscopy, 1 µl of cells and phage mix was spotted on an agarose pad (1.2% in MQ of UltraPure™ Agarose, Invitrogen) and directly imaged.

### TIRF microscopy acquisition and data evaluation

TIRF microscopy was performed using a Nikon Eclipse Ti2 inverted microscope equipped with an ILAS 2 TIRF module (Gataca Systems) and a TIRF 100x/1.49 oil objective. Samples were excited at 50 ms exposure for 14 frames using a 488 nm laser (200 mW) at 80% and emission was recovered via a quad TRIF filter cube (emission: 502-549 nm). The second frame in the fluorescent channel of the acquired TIRF microscopy images was denoised using the Nikon software package Denoise.ai. Subsequently, fluorescent maxima of ZorB, C or D translational fusions to mNeongreen were detected using MicrobeJ^57^ run in Fiji^58^. Statistics were calculated in Prism GraphPad 9^59^ by applying the in-built analyses of unpaired t-tests or one-way ANOVA.

### Bioinformatic analyses of ZorA motor and tail lengths

PADLOC v1.1.0^8,28^ with PADLOC-DB v1.4.0 was used to identify Zorya systems in RefSeq v209^60^ bacterial and archaeal genomes. Of the systems identified, we excluded those containing pseudogenes or more than one copy of each Zorya gene, or systems with non-canonical gene arrangements (Zorya genes are typically on the same strand, in type-specific conserved gene orders, e.g. ZorABCD, ZorABE, or ZorGABF for types I–III, respectively). To reduce redundancy due to highly related genome sequences in the RefSeq database, we then selected representative Zorya systems by clustering the sequences (using MMseqs2 v14.7e284^29^ with options: --min-seq-id 0.3 --coverage 0.8) of the proteins encoded by the three adjacent open reading frames on either side of each Zorya system, then randomly selected one system for each distinct set of these flanking genes (i.e. unique genetic context of the Zorya system). The ZorA and ZorB sequences from the representative Zorya systems were then clustered using MMseqs2 with options: --min-seq-id 0.3 - -coverage 0.95. Structures were predicted for one representative of each ZorA and ZorB cluster using ColabFold v1.5.2^42,61,62^ with options: --num-recycle 3 --num-models 1 --model-type auto -- amber --use-gpu-relax. Structures predictions were run as ZorA_5_ZorB_2_ multimers. The resulting structures were inspected manually (using PyMOL v2.5.4^47^) to identify the start of the ZorA tail. The rest of the sequences in each cluster were aligned to the representative sequence using MUSCLE v5.1^63^ using the Parallel Perturbed ProbCons algorithm (default) or the Super5 algorithm if the cluster contained more than 100 sequences. The start of the tail for the representative was used to infer the start of the tail for each other protein in the respective alignment.

### Figure preparation

Structural figures were prepared using ChimeraX^43^, PyMOL^47^, Prism GraphPad9 or GraphPad Prisim10^59^ and Adobe Illustrator^64^. The ion permeation pathway show in ZorAB was analyzed using MOLEonline^65^. The Hydrophobicity and polarity of the ZorAB tail was calculated using MOLEonline^65^. The electrostatic potential maps were calculated using the APBS^66^ electrostatic Plugin integrated inside Pymol.

## Data and Code Availability

Atomic coordinates for ZorAB WT, ZorA^E86A/E89A^ZorB, ZorA^Δ359–592^ZorB and ZorA^Δ435–729^ZorB were deposited in the Protein Data Bank (PDB) under accession codes 8QYD, 8QYH, 8QYK, 8QYY, respectively. The corresponding electrostatic potential maps were deposited in the Electron Microscopy Data Bank (EMDB) under accession codes EMD-18751, EMD-18754, EMD-18756, EMD-18766, respectively. The local refinement map of ZorB PGBD in ZorAB WT were deposited in the EMDB under accession codes EMD-18752. Atomic coordinates for ZorC were deposited in the PDB under accession codes PDB: 8R68. The corresponding electrostatic potential maps was deposited in the EMDB under accession codes EMDB: EMD-18848. Atomic coordinates for ZorD apo form and its complex with ATP-γ-S were deposited in the PDB under accession codes PDB: 8QY7 and 8QYC, respectively. The corresponding electrostatic potential maps were deposited in the EMDB under accession codes EMD-18747 and EMD-18750. The mass spectrometry proteomics data have been deposited to the ProteomeXchange Consortium via the PRIDE^67^ partner repository with the dataset identifier PXD047450. Username. Password: D7vNT520

## Acknowledgements

The Novo Nordisk Foundation Center for Protein Research is supported financially by the Novo Nordisk Foundation (NNF14CC0001). N.M.I.T. acknowledges support from NNF Hallas-Møller Emerging Investigator grant (NNF17OC0031006) and an NNF Project grant (NNF21OC0071948). H.H. acknowledges support from Lundbeck Foundation postdoc R347-2020-2429. S.A.J. acknowledges support from the Health Research Council of New Zealand (Sir Charles Hercus Fellowship) and from Bioprotection Aotearoa (Tertiary Education Commission, New Zealand). We also acknowledge the use of the New Zealand eScience Infrastructure (NeSI) high-performance computing facilities in this research. T.C.D.H. was supported by a University of Otago Doctoral Scholarship. M. E. acknowledges funding from the European Research Council (ERC) under the European Union’s Horizon 2020 research and innovation programme (grant agreement n ° 864971) and from the Max Planck Society as Max Planck Fellow. Y.W. acknowledges support from the National Key Research and Development Program of China (2021YFF1200404), the National Science Foundation of China (32371300), and computational resources from the Information Technology Center and State Key Lab of Computer-Aided Design (CAD) & Computer Graphics (CG) of Zhejiang University. M. L. N. lab was supported by the Novo Nordisk Foundation (NNF14CC0001), The Danish Council of Independent Research (grant agreement numbers 4002-00051, 4183-00322A and 8020-00220B), and The Danish Cancer Society (grant agreement R146-A9159-16-S2). We thank the Danish Cryo-EM Facility at the Core Facility for Integrated Microscopy (CFIM) at the University of Copenhagen and Tillmann Pape and Nicholas Heelund Sofos for support during data collection. We thank Blanca Lopez Mendez and Morten Ib Rasmussen for their support in Mass Spectrometry and Mass Photometry.

## Author contribution

N.M.I.T. and H.H. conceived the project. H.H., A.R-E., and N.R. and F.J.O.M. did molecular biology and mutagenesis. H.H. expressed, purified, optimized, prepared cryo-EM grids, collected cryo-EM data, and determined all the structures presented in this study. S.A.J. and T.C.D.H. performed phage infectivity, adsorption, and phage burst assays. S.A.J. performed cell survival, conjugation, and transformation assays. L. J. P. performed bioinformatic analyses. P.F.P. carried out and analyzed TIRF experiments and together with M. E. interpreted the localization studies. F.J.O.M. and N.R. assisted with protein purification. F.J.O.M., N.R., and H.H. optimized the nuclease and EMSA experiments. A.R-E. and V K.D.S. purified phage genomes. Y.W. and Y.Y. helped with ZorA tail structure modeling. H. H. prepared samples for mass spectrometry. I. A. H. and M. L. N. performed mass spectrometry and analyzed the data. H.H. prepared figures and wrote the first draft of the manuscript together with N.M.I.T. and S.A.J. with input from all the authors. This draft was then edited by M.E., P.F.P. and R.B., and all the other authors. All authors contributed to the revision of the manuscript.

## Extended Data Legends

**Extended Data Figure 1. *E. coli* Zorya type I protects against phage invasion but not bacterial conjugation or plasmid transformation.**

**a**, The impact of *Ec*ZorI on the uptake of plasmid DNA via conjugation from an *E. coli* donor strain, measured as the transconjugant frequency (number of transconjugants/total recipients). Four plasmids with different origins of replication (OriV) were tested (ColE1, RSF1010, pBBR1 and RK2), at the indicated donor to recipient cell ratios (D:R) for the matings. Data represent the mean of three replicates. **b**, The impact of *Ec*ZorI on the uptake of plasmid DNA via transformation. Chemically competent *E. coli* without (control; empty vector) or with *Ec*ZorI were transformed with plasmids possessing either ColE1 or pBBR1 origins of replication. Data represent the mean of three replicates, with each replicate being a different batch of competent cells. **c**, Infection time courses for liquid cultures of *E. coli*, with and without *Ec*ZorI, infected at different multiplicities of infection (MOI) of phage Bas02 and Bas08. **d**, Phage titers at the end timepoint for each sample from the infection time courses (**c**), measured as EOP on indicator lawns of *E. coli* either without (control) or with *Ec*ZorI. LOD: Limit of detection.

**Extended Data Figure 2. Cryo-EM dataset processing results and resolution of *Ec*ZorAB.**

**a**, A representative SDS gel of the purified *Ec*ZorAB complex. **b**, An EM image of the *Ec*ZorAB sample under the cryogenic conditions. **c**, Cryo-EM density map of *Ec*ZorAB colored by local resolution (in Å) estimated in cryoSPARC with gold standard (0.143) Fourier Shell Correlation (GSFSC) curves. **d**, Cryo-EM map of *Ec*ZorAB. **e-g**, Representative model segments of ZorA and non-residual molecules fitted into EM density, focusing on one of ZorA subunit’s TM1, and lipids found in the TMD of ZorA. **h**-**i**, Strategy of the local refinement of the ZorB PGBDs with a soft mask. **j**, A representative of a model segment of the ZorB PGBDs fitted into EM density map, focusing on the PGBD dimerized interface. **k**, Volcano plot analysis, visualizing ratio and significance of change between all proteins quantified by mass spectrometry in *E. coli* total lysates either transformed with p*Ec*ZorI plasmids or not (**Extended Data Table 2**). Significance was tested via two-tailed two-sample Student’s t-testing with permutation-based FDR control, ensuring a corrected p-value of < 0.01. n=4 technical replicates derived from n=3 culture replicates. **l**, Absolute copy-number analysis of Zorya proteins expressed in *E. coli*. Determined via comparison of molecular weight-adjusted label-free quantified protein abundance values from this study, to known copy-numbers reported by Schmidt et al.^56^, and establishing a “proteomic ruler” for conversion of measured abundance values to approximate copy-numbers (**Extended Data Table 2**). n=4 technical replicates derived from n=3 culture replicates.

**Extended Data Figure 3. *Ec*ZorA tail secondary structural prediction and a complete composite model of *Ec*ZorAB complex.**

**a**, Amino acids and secondary structural predictions (Psipred) of the *Ec*ZorA. The peptides found by mass spectrometry that covered ZorA protein are indicated as green lines about the amino acids. **b,** Top hits from an HHpred sequence homology search of the ZorA tail are shown. **c**, A composite model of *Ec*ZorAB with the ZorA tail folding into a pentameric super coiled-coil, with the helical pitch of the tail α-helix shown. **d**, Hydrophobicity and polarity of the inner surface of the ZorA tai calculated by MOLE*online*.

**Extended Data Figure 4. *Ec*ZorAB is a peptidoglycan binding rotary motor.**

**a**, Cartoon representation of the *Ec*ZorAB complex in an inactive state, with the ZorB dimerized interfaced highlighted. **b**, Topology diagrams of ZorB and isolated crystal structures of the flagellar stator unit MotB and PomB PGBDs, indicating a conserved folding architecture. **c**-**d**, The two disulfate bonds identified from ZorB PGBDs, with the EM map overlapped. **e**, Pull-down assay of the isolated *Ec*ZorAB complex with the purified peptidoglycan. **f**, Cartoon representation of the cryo-EM structure of the proton-driven flagellar stator unit MotAB from *Campylobacter jejuni* (*Cj*MotAB) in its inactive state, with the MotB plug motif highlighted. **g**, Cartoon representation of the cryo-EM structure of the sodium-driven flagellar stator unit PomAB from *Vibrio alginolyticus* (*Va*PomAB) in its inactive state. **h**, Cross-section view of the *Ec*ZorAB TMD, showing the surrounding residues of the two Asp26 from ZorB. **i**, Cross-section view of the *Cj*MotAB TMD, showing the surrounding residues of the two Asp22 from MotB. **j**, Cross-section view of *Va*PomAB TMD, showing the surrounding residues of the two Asp24 from PomB. The absence of the strictly conserved threonine residue on ZorA TM3 required for sodium ion binding, indicates that *Ec*ZorAB is a proton-driven stator unit. **m**, A representative of an SDS gel of the purified *Ec*ZorAB linker mutant complex (with ZorB residues 46-52 replaced by a GGGSGGS linker: *Ec*ZorAB linker mutant). **n**, An EM image of *Ec*ZorAB linker mutant sample under the cryogenic conditions. **o**, Representatives of the 2D classes of the *Ec*ZorAB linker mutant in comparison with that of the *Ec*ZorAB wild type, highlighting the flexibility of the ZorB PGBDs of the *Ec*ZorAB linker mutant. **p**, Low pass filter of the Cryo-EM density map of the *Ec*ZorAB linker mutant after nonuniform refinement. **q**, Transmembrane helix density of the *Ec*ZorAB linker mutant and that in the wild type *Ec*ZorAB.

**Extended Data Figure 5. Cryo-EM dataset processing and structures of the *Ec*ZorAB tail and Ca^2+^ binding site mutations.**

**A**, Mutations of the ZorA tail truncations indicated in the composite model of *Ec*ZorAB complex. **B**, Interaction between the beginning of the ZorA tail and the β-hairpin motif. **C**, Extra density found inside the tail from cryo-EM map, which was modeled as palmitic acid, with the amino acids involved in the interactions indicated. **d**, Structural comparison of the ZorA wild type (cyan) and ZorA Ca^2+^ binding site mutation (ZorA^E86A/E89A^, gray), the arrows highlight the changes from wild type to the mutant. **e**, Representative of the 2D classes of the *Ec*ZorAB ZorA tail complete deletion. **f**, Negative staining images of the *Ec*ZorAB wild type, ZorA tail middle deletion (ZorA^Δ359–592^), ZorA tail tip deletion (ZorAΔ^435–729^). **g**, The tail lengths of the *Ec*ZorAB wild type, ZorA tail middle deletion (ZorA^Δ359–592^), ZorA tail tip deletion (ZorA^Δ359–592^) as measured in (**f**). **h**-**j**, Cryo-EM maps and resolutions of ZorA mutants with gold standard (0.143) Fourier Shell Correlation (GSFSC) curves.

**Extended Data Figure 6. The effects of *Ec*Zorya mutations on *Ec*ZorI-mediated anti-phage defense and long ZorA tails are conserved amongst Zorya system types in diverse species.**

**a**, The effects of ZorA, ZorB, ZorC and ZorD mutations on *Ec*ZorI-mediated anti-phage defense, as measured using EOP assays with phages Bas02, Bas19 and Bas25. Data represent the mean of at least 3 replicates and are normalized to the control samples lacking *Ec*ZorI. **b**, The ZorA tail lengths found in different Zorya system types. Motor and tail lengths were determined by inspecting the predicted structures of several representative ZorA sequences, then inferring these lengths for the rest of the ZorA sequences through sequence alignment (methods). The reduce sequencing bias, unique Zorya systems encoded in RefSeq (v209) bacteria and archaea genomes were selected based on their distinct genomic context (methods).

**Extended Data Figure 7. Cryo-EM dataset processing results and functional investigation of *Ec*ZorC.**

**a**, Representative of the SDS gel of the purified ZorC wild type, ZorC^E400A^, ZorC^H443A^, ZorC^ΔCTD^ (deletion residues 487-560). **b**, Representatives of the 2D classes of the *Ec*ZorC. **c**, Unsharpened Cryo-EM map of *Ec*ZorC with gold standard (0.143) Fourier Shell Correlation (GSFSC) curves shown below. **d**, Local refinement of the *Ec*ZorC core domain with a soft mask, with the local resolution (in Å) estimated in cryoSPARC. **e**, Representative of a model and segments of the ZorC fitted into EM density map. **f**, Final model of *Ec*ZorC built from cryo-EM map. **g**, AlphaFold2-predicted ZorC model. **h**, Electrostatic distribution of *Ec*ZorC calculated from AlphaFold2-predicted model.

**Extended Data Figure 8. Cryo-EM dataset processing results and resolutions of *Ec*ZorD and *Ec*ZorD in complex with ATP-γ-S.**

**a**, Representative of the SDS gel of the purified ZorD wild type, ZorD_CTD_ (residues 503-1080), ZorD_NTD_ (residues 1-502), ZorD_CTD_^D730A/E731A^ and ZorD_CTD_^E651A^. Gel is representative of at least 3 replicates. **b**, Unsharpened cryo-EM map of the *Ec*ZorD apo from with gold standard (0.143) Fourier Shell Correlation (GSFSC) curves shown below. **c**, Local refinement of the *Ec*ZorD apo form with a soft mask. **d**, Cryo-EM map of *Ec*ZorD in complex with ATP-γ-S. **e**, Structural model of the *Ec*ZorD in complex with ATP-γ-S. **f**, Structural comparison of the *Ec*ZorD apo from (gray) and *Ec*ZorD in complex with ATP-γ-S (light purple); the arrows highlight the changes from apo form to the ligand-bound form. **g**, Zoomed-in view of the ATP-γ-S binding site, with the cryo-EM map overlayed on the ATP-γ-S.

**Extended Data Figure 9. Complementation experiment between *E. coli* and *P. aeruginosa* ZoryaI.**

**a**, Schematic representation of *Ec*ZorI, *Pa*ZorI and the constructs for *Pa*ZorCD or *Pa*ZorD complementation of *Ec*ZorI gene deletions. **b**, Anti-phage defense provided by the constructs in (**a**), as measured using EOP assays for phages Bas49, Bas52 and Bas57. Data represent the mean of at least 3 replicates and are normalized to the control samples lacking Zorya.

**Extended Data Figure 10. ZorD recruitment during phage invasion at increasing MOIs and mNeongreen localization upon phage exposure.**

**a**, The effects of the mNeongreen (mNG) fusions to *Ec*ZorI components on anti-phage defense, as measured using EOP assays for phages Bas02, Bas08, Bas19, Bas24, Bas25 and Bas58. Data represent the mean of at least 3 replicates and are normalized to the control samples lacking *Ec*ZorI. The boxed constructs (ZorB C-terminal mNG fusion: ZorB-mNG; ZorC N-terminal mNG fusion: mNG-ZorB; ZorD C-terminal mNG fusion: ZorD-mNG) were used for subsequent microscopy experiments. **b**, Exemplary denoised TIRF and brightfield microscopy pictures of mNeongreen expression driven by the *Ec*ZorI native promoter (p-mNG) either untreated, exposed to T4 or Bas24 at an MOI of 5 for 30 min. Scale bar 2 µm. **c**, **e**, Exemplary denoised TIRF microscopy pictures of ZorD-mNG either untreated or exposed to increasing Bas24 or T4 MOIs of 1, 5, or 50 for 30 min. **d**, **f**, Statistical comparison of ZorD-mNG maxima between untreated and conditions stated in **c**, **e**. Means derive from at least three independent biological replicates. Scale bar 2 µm. Our data in **e** and **f** showed phage T4 infection did not result in a dose-dependent ZorD-mNG response.

**Extended Data Figure 11. Structural prediction of the representative ZorAB complexes form different Zorya system types.**

**a**, The predicted dimerized ZorB PGBDs. **b**, The predicted ZorAB transmembrane motor complex. One ZorA subunit is highlighted.

**Extended Data Fig. 12. Proposed ZorA tail untwisting and phage DNA ‘reeling in’ mechanism in the activated Zorya defense system.**

**a,** An inactive ZorAB embedded in the inner membrane. **b,** Phage invasion triggers ZorAB activation. The rotation of ZorA and its long intracellular tail around ZorB causes untwisting of the ZorA tail, which would recruit ZorC and ZorD**. c,** Reeling in of phage DNA around the long ZorA tail in the activated Zorya defense system. Legend Discussion: The ‘reeling in’ mechanism would greatly enhance phage genome localization and sequester it from interactions required for host infection. A typical double-stranded DNA phage genome is 10s of μm long, and would form a random coil with a radius of gyration similar to the size of the entire cell in the absence of constraints – thus negating any advantage of a localized nuclease defense response at the site of entry. Binding to multiple Zor complexes, perhaps via ZorC and/or D might contribute to localizing phage DNA to the entry site. Unless and until tail rotation is resisted, an almost inevitable consequence of tail rotation combined with DNA binding is that the DNA will wind around the tail like wire on a reel. If ‘reeling in’ of the phage DNA were an essential feature of the Zorya defense mechanism, this would explain the need for both rotation and a long ZorA tail. The length of the ZorA tail is very similar to that of a typical phage capsid into which an entire phage genome can (only just) be tightly packed, indicating that a single Zor complex may be sufficient to capture an entire genome. Rough calculations indicate that 100s of turns would be required to wind a 60 kb Bas24 genome onto a 70 nm tail, allowing the rotary ZorAB motor cumulatively to supply the necessary energy to wind and compact the phage DNA. Reeling would also tighten any loops that might form between DNA sites bound to different ZorA tails, removing any freedom for further ZorA rotation - as required by the model of activation via untwisting of the ZorA tail.

## Extended Data Tables

**Extended Data Table 1. Cryo-EM data collection, refinement and validation statistics.**

**Extended Data Table 2. Mass Spectrometry analysis of the *E.coli* lysates with or without expression of the Zorya proteins.**

## Extended Data Videos

**Extended Data Video 1 and Video 2.**

**Exemplary deconvolved TIRF microscopy videos (1800 ms) of ZorB C-terminal fusion with mNeongreen.**

*E. coli* expressing Zorya untreated with phage (**Video 1**) or exposed to Bas24 (**Video 2**) at an MOI of 5 for 30 min. The ZorAB complexes were predominantly membrane-localized and exhibited low diffusibility, irrespective of the absence or presence of phage (**Video 1)**. Upon phage infection, the number of ZorAB foci increased (**Video 2**). Scale bar 2 µm.

**Extended Data Video 3 and Video 4.**

**Exemplary deconvolved TIRF microscopy videos (1800 ms) of ZorD C-terminal fusion with mNeongreen**

*E. coli* expressing Zorya untreated with phage (**Video 3**) or exposed to Bas24 (**Video 4**) at an MOI of 5 for 30 min. ZorD freely diffused in the cytoplasm in the absence of phage (**Video 3)**. Upon phage infection, the number of ZorD foci increased and became static (**Video 4**). Scale bar 2 µm.

