## Supplemental figures for "Structure and mechanism of Zorya anti-phage defense system"

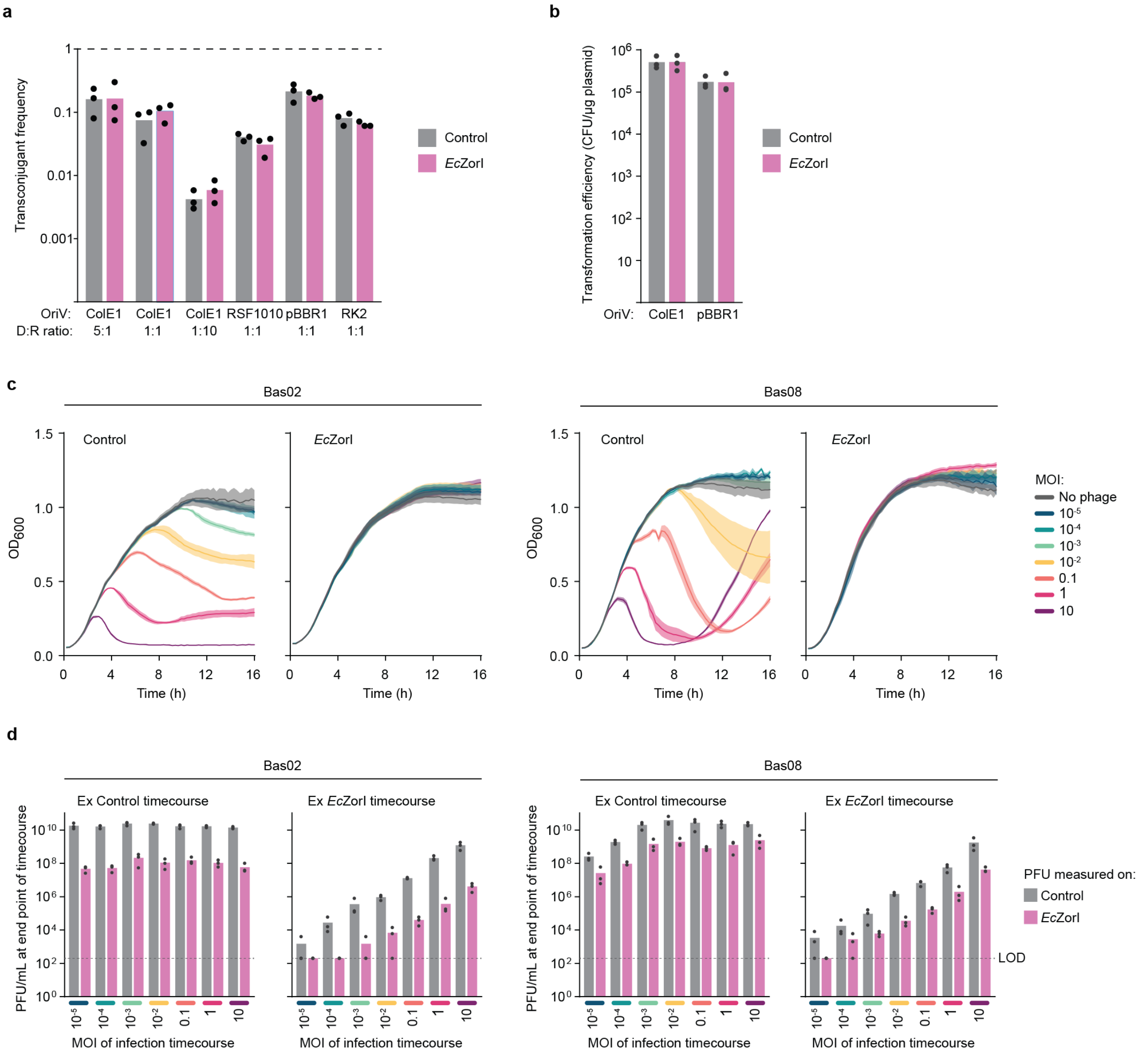


**Extended Data Figure 1. *E. coli* Zorya type I protects against phage invasion but not bacterial conjugation or plasmid transformation.**

**a**, The impact of *Ec*ZorI on the uptake of plasmid DNA via conjugation from an *E. coli* donor strain, measured as the transconjugant frequency (number of transconjugants/total recipients). Four plasmids with different origins of replication (OriV) were tested (ColE1, RSF1010, pBBR1 and RK2), at the indicated donor to recipient cell ratios (D:R) for the matings. Data represent the mean of three replicates. **b**, The impact of *Ec*ZorI on the uptake of plasmid DNA via transformation. Chemically competent *E. coli* without (control; empty vector) or with *Ec*ZorI were transformed with plasmids possessing either ColE1 or pBBR1 origins of replication. Data represent the mean of three replicates, with each replicate being a different batch of competent cells. **c**, Infection time courses for liquid cultures of *E. coli*, with and without *Ec*ZorI, infected at different multiplicities of infection (MOI) of phage Bas02 and Bas08. **d**, Phage titers at the end timepoint for each sample from the infection time courses (**c**), measured as EOP on indicator lawns of *E. coli* either without (control) or with *Ec*ZorI. LOD: Limit of detection.

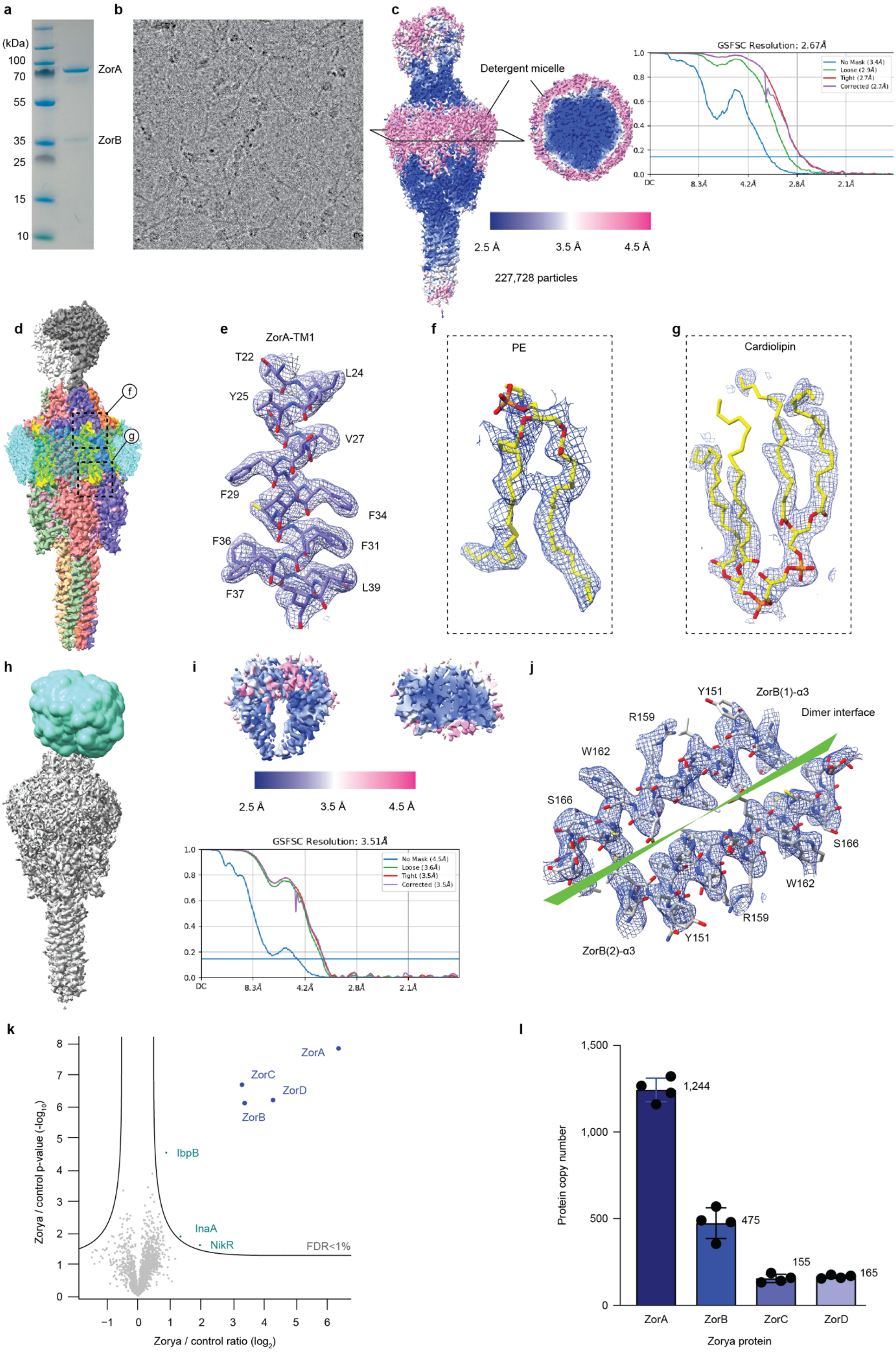
a
(kDa)
100
70
55
35
25
15
10
ZorA
ZorB
b
c
Detergent micelle
GSFSC Resolution: 2.67Å
No Mask (3.4Å)
Loose (2.9Å)
Tight (2.7Å)
Corrected (2.7Å)
2.5 Å
3.5 Å
4.5 Å
227,728 particles
d
e
ZorA-TM1
T22
L24
Y25
V27
F29
F34
F36
F31
F37
L39
f
PE
g
Cardiolipin
h
i
2.5 Å
3.5 Å
4.5 Å
GSFSC Resolution: 3.51Å
No Mask (4.5Å)
Loose (3.6Å)
Tight (3.5Å)
Corrected (3.5Å)
j
Y151
ZorB(1)-α3
Dimer interface
R159
W162
S166
S166
W162
R159
Y151
ZorB(2)-α3
k
ZorA
ZorC
ZorD
ZorB
IbpB
InaA
NikR
FDR<1%
Zorya / control p-value (-log10)
Zorya / control ratio (log2)
l
1,500
1,000
500
0
1,244
475
155
165
Protein copy number
ZorA
ZorB
ZorC
ZorD
Zorya protein

**Extended Data Figure 2. Cryo-EM dataset processing results and resolution of *Ec*ZorAB.** **a**, A representative SDS gel of the purified *Ec*ZorAB complex. **b**, An EM image of the *Ec*ZorAB sample under the cryogenic conditions. **c**, Cryo-EM density map of *Ec*ZorAB colored by local resolution (in Å) estimated in cryoSPARC with gold standard (0.143) Fourier Shell Correlation (GSFSC) curves. **d**, Cryo-EM map of *Ec*ZorAB. **e-g**, Representative model segments of ZorA and non-residual molecules fitted into EM density, focusing on one of ZorA subunit's TM1, and lipids found in the TMD of ZorA. **h**-**i**, Strategy of the local refinement of the ZorB PGBDs with a soft mask. **j**, A representative of a model segment of the ZorB PGBDs fitted into EM density map, focusing on the PGBD dimerized interface. **k**, Volcano plot analysis, visualizing ratio and significance of change between all proteins quantified by mass spectrometry in *E. coli* total lysates either transformed with p*Ec*ZorI plasmids or not (**Extended Data Table 2**). Significance was tested via two-tailed two-sample Student's t-testing with permutation-based FDR control, ensuring a corrected p-value of < 0.01. n=4 technical replicates derived from n=3 culture replicates. **l**, Absolute copy-number analysis of Zorya proteins expressed in *E. coli*. Determined via comparison of molecular weight-adjusted label-free quantified protein abundance values from this study, to known copy-numbers reported by Schmidt et al.[56], and establishing a "proteomic ruler" for conversion of measured abundance values to approximate copy-numbers (**Extended Data Table 2**). n=4 technical replicates derived from n=3 culture replicates.

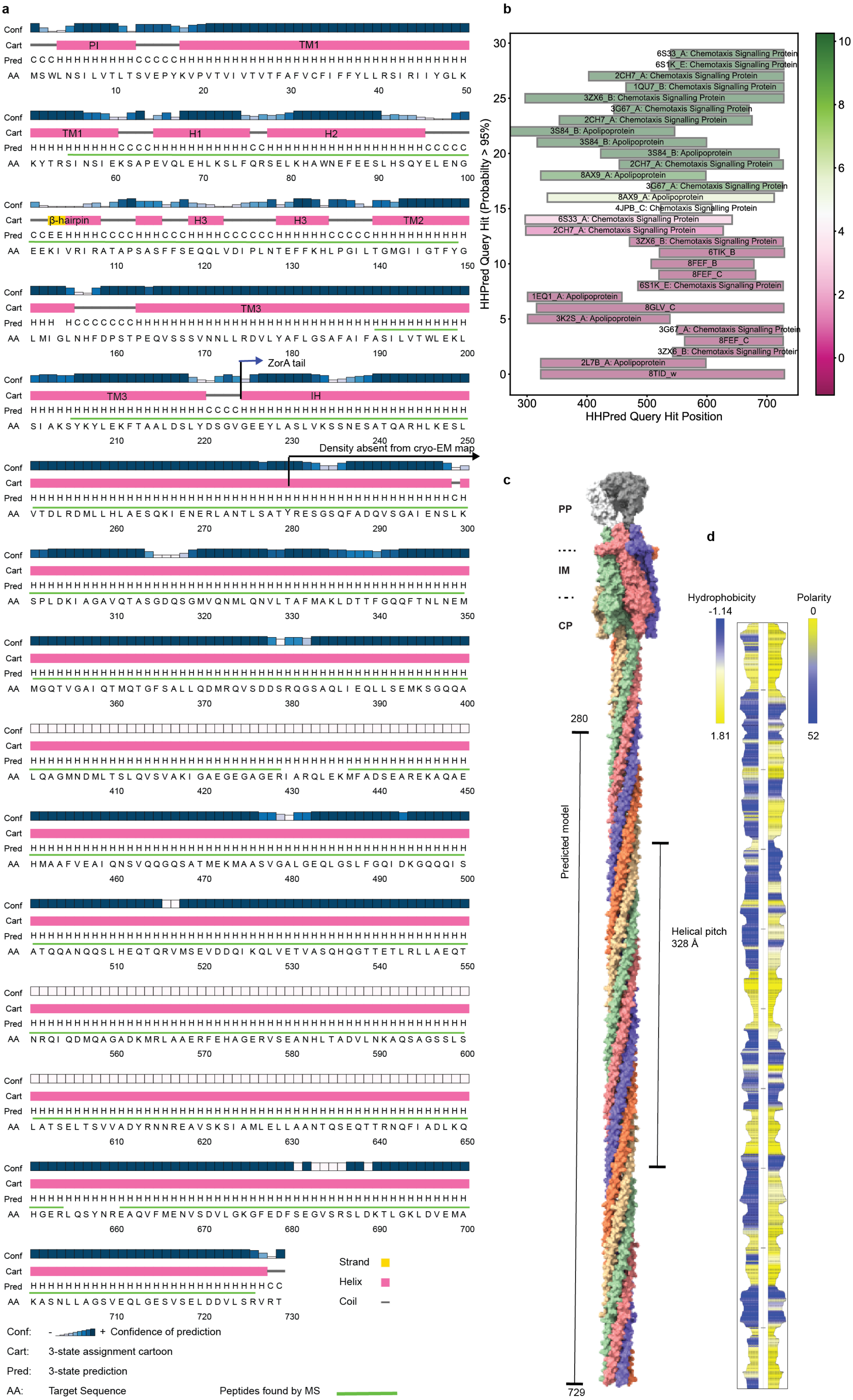
a
Conf
Cart
Pred
AA
PI
TM1
H1
H2
β-hairpin
H3
TM2
TM3
IH
ZorA tail
Density absent from cryo-EM map
Strand
Helix
Coil
Conf: - + Confidence of prediction
Cart: 3-state assignment cartoon
Pred: 3-state prediction
AA: Target Sequence
Peptides found by MS
b
HHPred Query Hit (Probability > 95%)
HHPred Query Hit Position
-log(E-value)
6S33_A: Chemotaxis Signalling Protein
6S1K_E: Chemotaxis Signalling Protein
2CH7_A: Chemotaxis Signalling Protein
1QU7_B: Chemotaxis Signalling Protein
3ZX6_B: Chemotaxis Signalling Protein
3G67_A: Chemotaxis Signalling Protein
3S84_B: Apolipoprotein
8AX9_A: Apolipoprotein
4JPB_C: Chemotaxis Signalling Protein
6TIK_B
8FEF_B
8FEF_C
1EQ1_A: Apolipoprotein
8GLV_C
3K2S_A: Apolipoprotein
2L7B_A: Apolipoprotein
8TID_w
c
PP
IM
CP
280
Predicted model
Helical pitch
328 Å
729
d
Hydrophobicity
-1.14
1.81
Polarity
0
52

**Extended Data Figure 3. *Ec*ZorA tail secondary structural prediction and a complete composite model of *Ec*ZorAB complex.**

**a**, Amino acids and secondary structural predictions (Psipred) of the *Ec*ZorA. The peptides found by mass spectrometry that covered ZorA protein are indicated as green lines about the amino acids.

**b,** Top hits from an HHpred sequence homology search of the ZorA tail are shown. **c**, A composite model of *Ec*ZorAB with the ZorA tail folding into a pentameric super coiled-coil, with the helical pitch of the tail $\alpha$-helix shown. **d**, Hydrophobicity and polarity of the inner surface of the ZorA tai calculated by MOLE*online*.

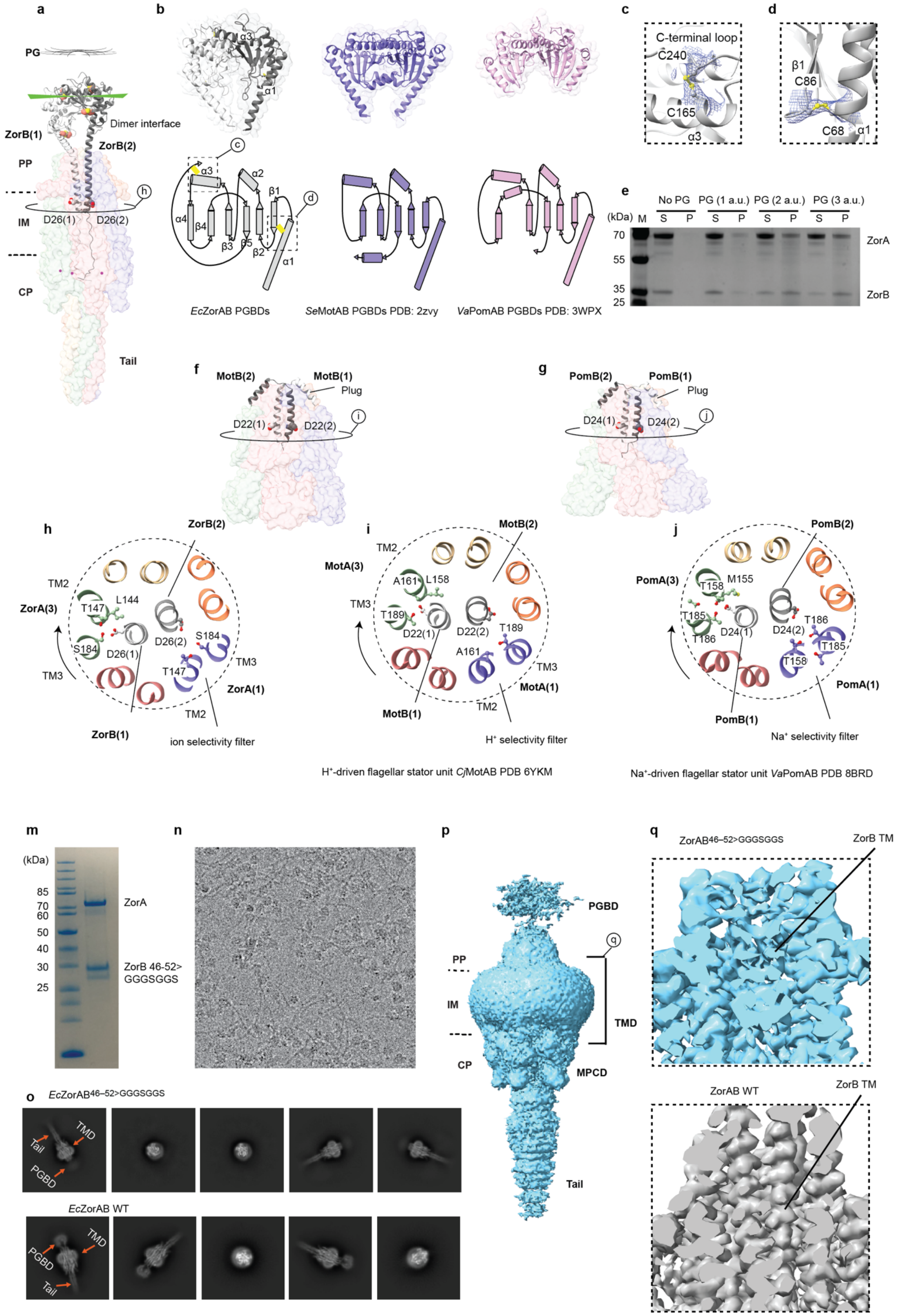
a
PG
Dimer interface
ZorB(1)
ZorB(2)
PP
IM
CP
D26(1)
D26(2)
Tail
b
EcZorAB PGBDs
SeMotAB PGBDs PDB: 2zvy
VaPomAB PGBDs PDB: 3WPX
c
C-terminal loop
C240
C165
α3
d
β1
C86
C68
α1
e
No PG
PG (1 a.u.)
PG (2 a.u.)
PG (3 a.u.)
(kDa) M
S P
70
55
35
25
ZorA
ZorB
f
MotB(2)
MotB(1)
Plug
D22(1)
D22(2)
g
PomB(2)
PomB(1)
Plug
D24(1)
D24(2)
h
ZorB(2)
ZorA(3)
TM2
TM3
T147
L144
S184
D26(1)
D26(2)
ZorA(1)
ZorB(1)
ion selectivity filter
i
MotB(2)
MotA(3)
A161
L158
T189
D22(1)
D22(2)
MotA(1)
MotB(1)
H⁺ selectivity filter
H⁺-driven flagellar stator unit CjMotAB PDB 6YKM
j
PomB(2)
PomA(3)
T158
M155
T185
T186
D24(1)
D24(2)
PomA(1)
PomB(1)
Na⁺ selectivity filter
Na⁺-driven flagellar stator unit VaPomAB PDB 8BRD
m
(kDa)
85
70
60
50
40
30
25
ZorA
ZorB 46-52>
GGGSGGS
n
o
EcZorAB46–52>GGGSGGS
TMD
Tail
PGBD
EcZorAB WT
p
PGBD
PP
IM
CP
TMD
MPCD
Tail
q
ZorAB46–52>GGGSGGS
ZorB TM
ZorAB WT

**Extended Data Figure 4. *Ec*ZorAB is a peptidoglycan binding rotary motor.** **a**, Cartoon representation of the *Ec*ZorAB complex in an inactive state, with the ZorB dimerized interfaced highlighted. **b**, Topology diagrams of ZorB and isolated crystal structures of the flagellar stator unit MotB and PomB PGBDs, indicating a conserved folding architecture. **c**-**d**, The two disulfate bonds identified from ZorB PGBDs, with the EM map overlapped. **e**, Pull-down assay of the isolated *Ec*ZorAB complex with the purified peptidoglycan. **f**, Cartoon representation of the cryo-EM structure of the proton-driven flagellar stator unit MotAB from *Campylobacter jejuni* (*Cj*MotAB) in its inactive state, with the MotB plug motif highlighted. **g**, Cartoon representation of the cryo-EM structure of the sodium-driven flagellar stator unit PomAB from *Vibrio alginolyticus* (*Va*PomAB) in its inactive state. **h**, Cross-section view of the *Ec*ZorAB TMD, showing the surrounding residues of the two Asp26 from ZorB. **i**, Cross-section view of the *Cj*MotAB TMD, showing the surrounding residues of the two Asp22 from MotB. **j**, Cross-section view of *Va*PomAB TMD, showing the surrounding residues of the two Asp24 from PomB. The absence of the strictly conserved threonine residue on ZorA TM3 required for sodium ion binding, indicates that *Ec*ZorAB is a proton-driven stator unit. **m**, A representative of an SDS gel of the purified *Ec*ZorAB linker mutant complex (with ZorB residues 46-52 replaced by a GGGSGGS linker: *Ec*ZorAB linker mutant). **n**, An EM image of *Ec*ZorAB linker mutant sample under the cryogenic conditions. **o**, Representatives of the 2D classes of the *Ec*ZorAB linker mutant in comparison with that of the *Ec*ZorAB wild type, highlighting the flexibility of the ZorB PGBDs of the *Ec*ZorAB linker mutant. **p**, Low pass filter of the Cryo-EM density map of the *Ec*ZorAB linker mutant after nonuniform refinement. **q**, Transmembrane helix density of the *Ec*ZorAB linker mutant and that in the wild type *Ec*ZorAB.

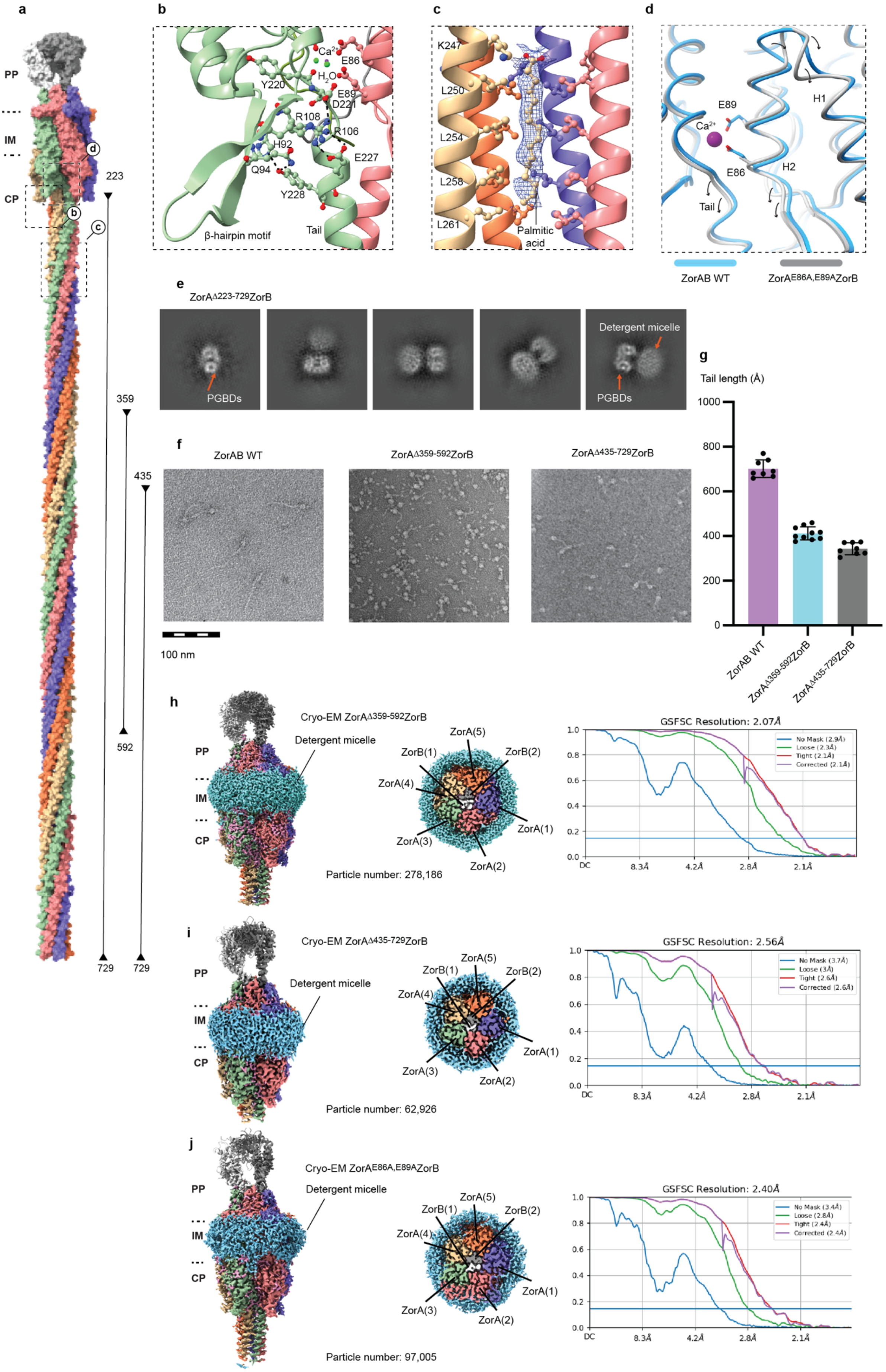
a
b
c
d
PP
IM
CP
223
359
435
592
729
729
Ca2+
E86
H2O
Y220
E89
D221
R108
R106
H92
E227
Q94
Y228
β-hairpin motif
Tail
K247
L250
L254
L258
L261
Palmitic acid
E89
Ca2+
H1
H2
E86
Tail
ZorAB WT
ZorAE86A,E89AZorB
e
ZorAΔ223-729ZorB
Detergent micelle
PGBDs
PGBDs
g
Tail length (Å)
1000
800
600
400
200
0
ZorAB WT
ZorAΔ359-592ZorB
ZorAΔ435-729ZorB
f
ZorAB WT
ZorAΔ359-592ZorB
ZorAΔ435-729ZorB
100 nm
h
Cryo-EM ZorAΔ359-592ZorB
Detergent micelle
PP
IM
CP
ZorA(5)
ZorB(1)
ZorB(2)
ZorA(4)
ZorA(1)
ZorA(3)
ZorA(2)
Particle number: 278,186
GSFSC Resolution: 2.07Å
No Mask (2.9Å)
Loose (2.3Å)
Tight (2.1Å)
Corrected (2.1Å)
DC
8.3Å
4.2Å
2.8Å
2.1Å
i
Cryo-EM ZorAΔ435-729ZorB
Detergent micelle
ZorA(5)
ZorB(1)
ZorB(2)
ZorA(4)
ZorA(1)
ZorA(3)
ZorA(2)
Particle number: 62,926
GSFSC Resolution: 2.56Å
No Mask (3.7Å)
Loose (3Å)
Tight (2.6Å)
Corrected (2.6Å)
j
Cryo-EM ZorAE86A,E89AZorB
Detergent micelle
ZorA(5)
ZorB(1)
ZorB(2)
ZorA(4)
ZorA(1)
ZorA(3)
ZorA(2)
Particle number: 97,005
GSFSC Resolution: 2.40Å
No Mask (3.4Å)
Loose (2.8Å)
Tight (2.4Å)
Corrected (2.4Å)

**Extended Data Figure 5. Cryo-EM dataset processing and structures of the *Ec*ZorAB tail and $Ca^{2+}$ binding site mutations.**

**A**, Mutations of the ZorA tail truncations indicated in the composite model of *Ec*ZorAB complex. **B**, Interaction between the beginning of the ZorA tail and the β-hairpin motif. **C**, Extra density found inside the tail from cryo-EM map, which was modeled as palmitic acid, with the amino acids involved in the interactions indicated. **d**, Structural comparison of the ZorA wild type (cyan) and ZorA $Ca^{2+}$ binding site mutation (ZorA$^{E86A/E89A}$, gray), the arrows highlight the changes from wild type to the mutant. **e**, Representative of the 2D classes of the *Ec*ZorAB ZorA tail complete deletion. **f**, Negative staining images of the *Ec*ZorAB wild type, ZorA tail middle deletion (ZorA$^{\Delta359\text{-}592}$), ZorA tail tip deletion (ZorAΔ$^{435\text{-}729}$). **g**, The tail lengths of the *Ec*ZorAB wild type, ZorA tail middle deletion (ZorA$^{\Delta359\text{-}592}$), ZorA tail tip deletion (ZorA$^{\Delta359\text{-}592}$) as measured in (**f**). **h**-**j**, Cryo-EM maps and resolutions of ZorA mutants with gold standard (0.143) Fourier Shell Correlation (GSFSC) curves.

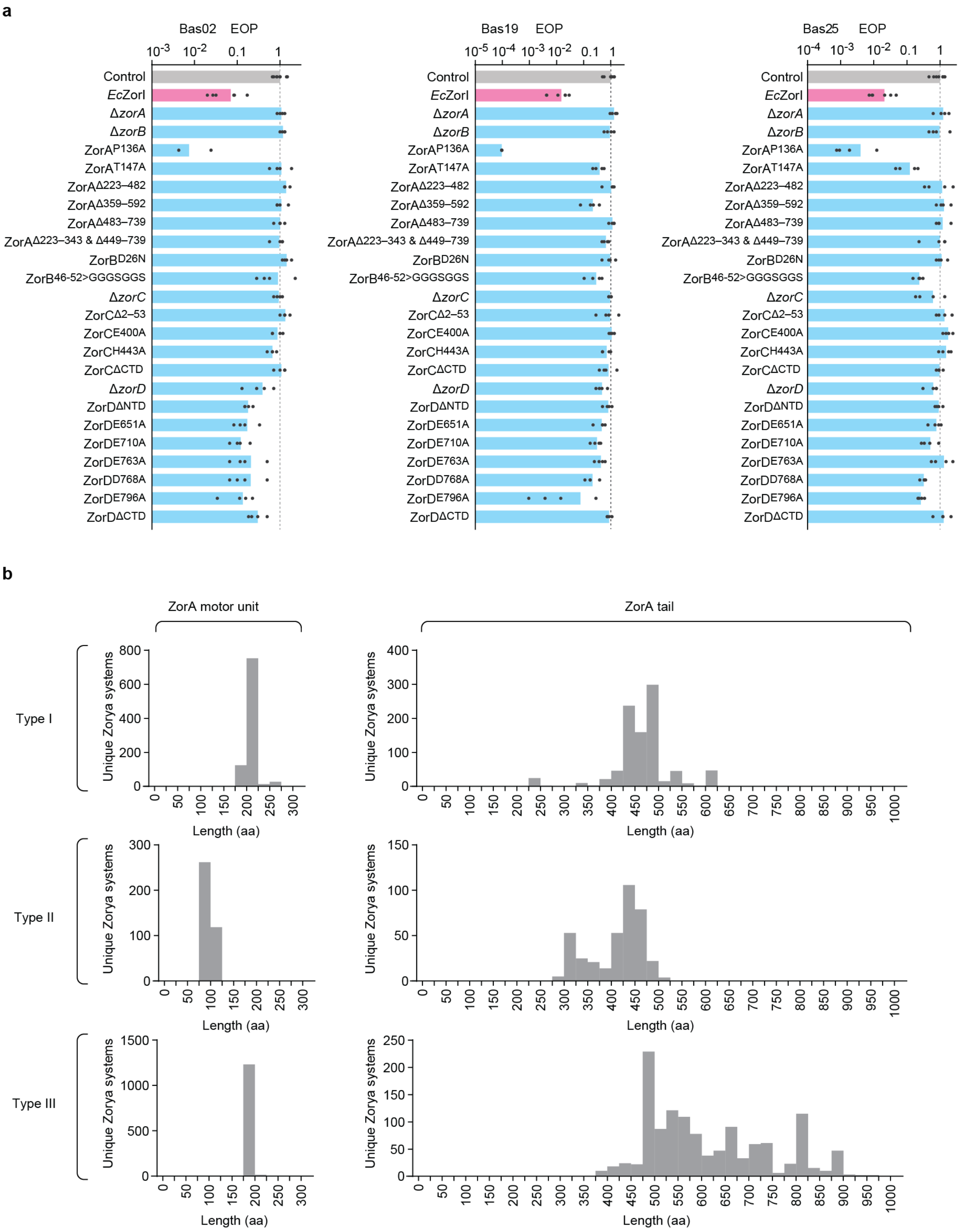
a
Bas02 EOP
Bas19 EOP
Bas25 EOP
Control
EcZorI
ΔzorA
ΔzorB
ZorAP136A
ZorAT147A
ZorAΔ223–482
ZorAΔ359–592
ZorAΔ483–739
ZorAΔ223–343 & Δ449–739
ZorBD26N
ZorB46-52>GGGSGGS
ΔzorC
ZorCΔ2–53
ZorCE400A
ZorCH443A
ZorCΔCTD
ΔzorD
ZorDΔNTD
ZorDE651A
ZorDE710A
ZorDE763A
ZorDD768A
ZorDE796A
ZorDΔCTD
b
ZorA motor unit
ZorA tail
Type I
Type II
Type III
Unique Zorya systems
Length (aa)

**Extended Data Figure 6. The effects of *Ec*Zorya mutations on *Ec*ZorI-mediated anti-phage defense and long ZorA tails are conserved amongst Zorya system types in diverse species.**

**a**, The effects of ZorA, ZorB, ZorC and ZorD mutations on *Ec*ZorI-mediated anti-phage defense, as measured using EOP assays with phages Bas02, Bas19 and Bas25. Data represent the mean of at least 3 replicates and are normalized to the control samples lacking *Ec*ZorI. **b**, The ZorA tail lengths found in different Zorya system types. Motor and tail lengths were determined by inspecting the predicted structures of several representative ZorA sequences, then inferring these lengths for the rest of the ZorA sequences through sequence alignment (methods). The reduce sequencing bias, unique Zorya systems encoded in RefSeq (v209) bacteria and archaea genomes were selected based on their distinct genomic context (methods).

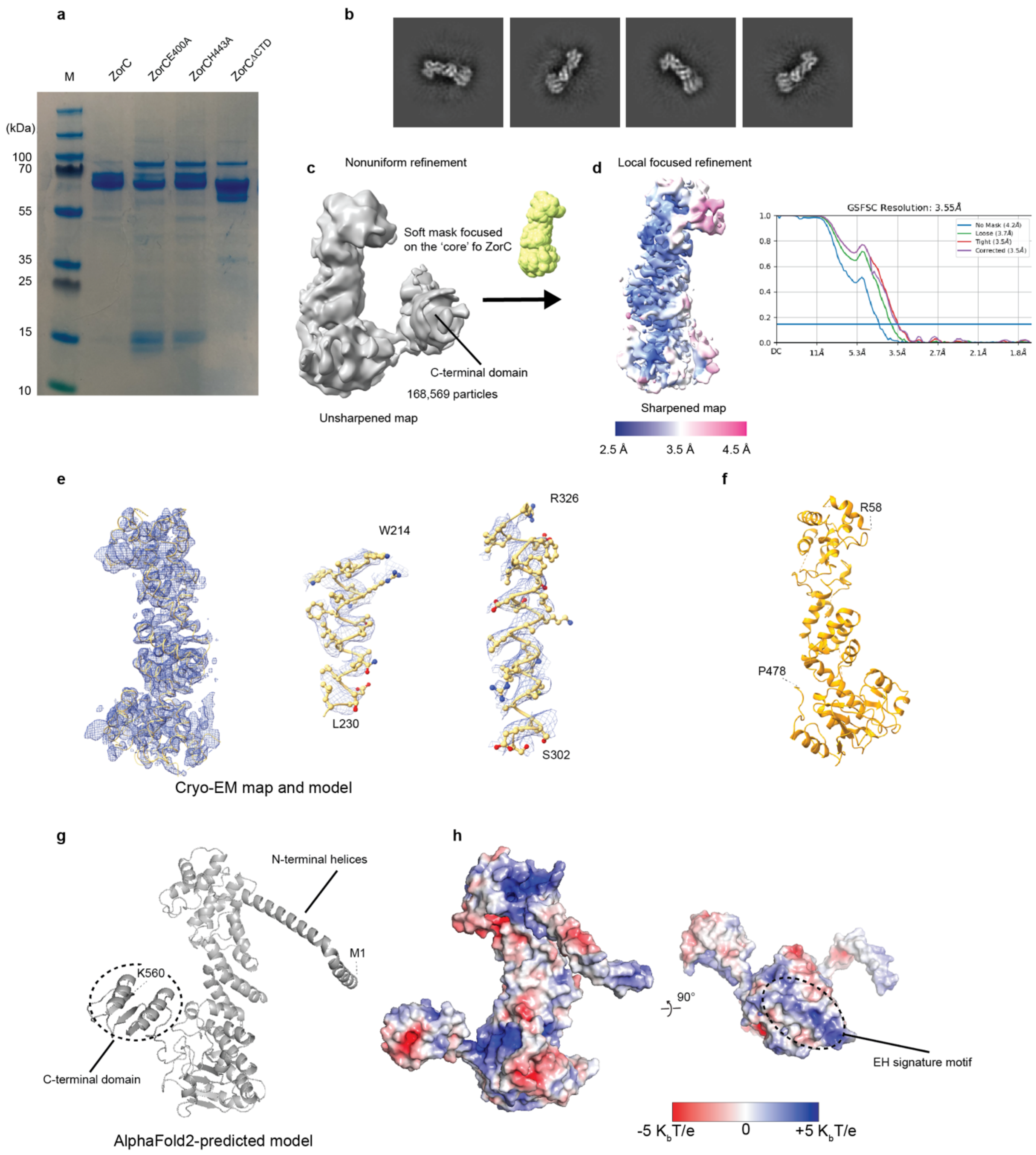


**Extended Data Figure 7. Cryo-EM dataset processing results and functional investigation of *Ec*ZorC.**

**a**, Representative of the SDS gel of the purified ZorC wild type, ZorC$^{E400A}$, ZorC$^{H443A}$, ZorC$^{\Delta CTD}$ (deletion residues 487-560). **b**, Representatives of the 2D classes of the *Ec*ZorC. **c**, Unsharpened Cryo-EM map of *Ec*ZorC with gold standard (0.143) Fourier Shell Correlation (GSFSC) curves shown below. **d**, Local refinement of the *Ec*ZorC core domain with a soft mask, with the local resolution (in Å) estimated in cryoSPARC. **e**, Representative of a model and segments of the ZorC fitted into EM density map. **f**, Final model of *Ec*ZorC built from cryo-EM map. **g**, AlphaFold2-predicted ZorC model. **h**, Electrostatic distribution of *Ec*ZorC calculated from AlphaFold2-predicted model.

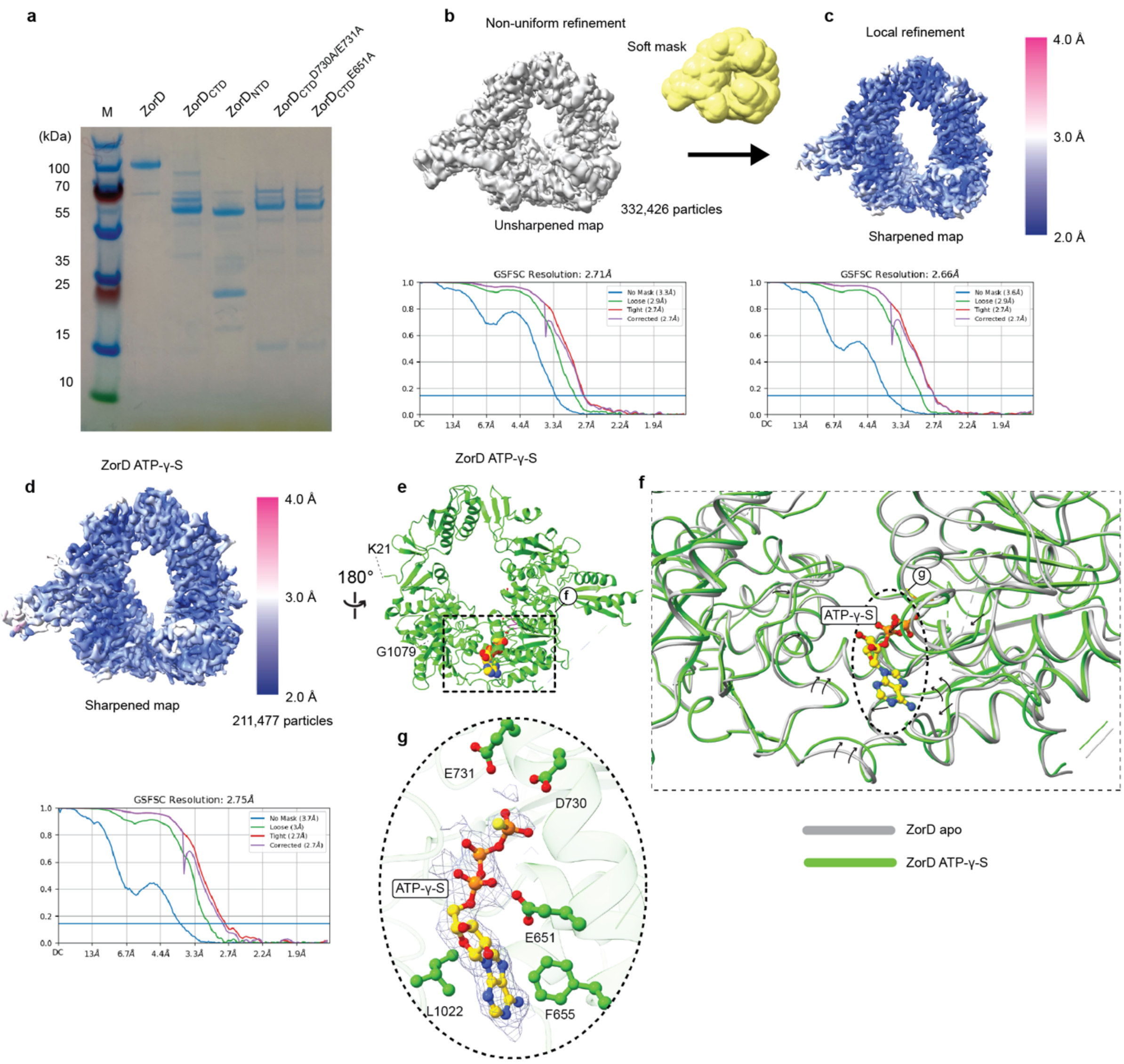


**Extended Data Figure 8. Cryo-EM dataset processing results and resolutions of *Ec*ZorD and *Ec*ZorD in complex with ATP-γ-S.**

**a**, Representative of the SDS gel of the purified ZorD wild type, ZorD$_{CTD}$ (residues 503-1080), ZorD$_{NTD}$ (residues 1-502), ZorD$_{CTD}^{D730A/E731A}$ and ZorD$_{CTD}^{E651A}$. Gel is representative of at least 3 replicates. **b**, Unsharpened cryo-EM map of the *Ec*ZorD apo from with gold standard (0.143) Fourier Shell Correlation (GSFSC) curves shown below. **c**, Local refinement of the *Ec*ZorD apo form with a soft mask. **d**, Cryo-EM map of *Ec*ZorD in complex with ATP-γ-S. **e**, Structural model of the *Ec*ZorD in complex with ATP-γ-S. **f**, Structural comparison of the *Ec*ZorD apo from (gray) and *Ec*ZorD in complex with ATP-γ-S (light purple); the arrows highlight the changes from apo form to the ligand-bound form. **g**, Zoomed-in view of the ATP-γ-S binding site, with the cryo-EM map overlayed on the ATP-γ-S.

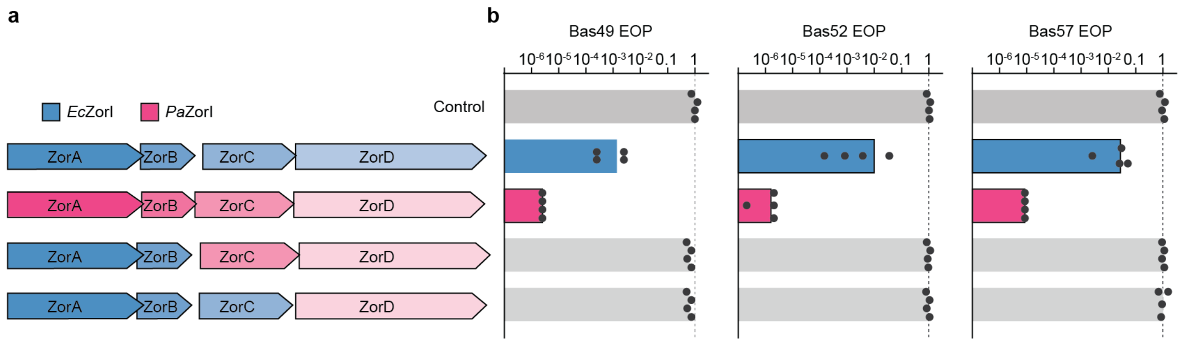


**Extended Data Figure 9. Complementation experiment between *E. coli* and *P. aeruginosa* ZoryaI.**
**a**, Schematic representation of *Ec*ZorI, *Pa*ZorI and the constructs for *Pa*ZorCD or *Pa*ZorD complementation of *Ec*ZorI gene deletions. **b**, Anti-phage defense provided by the constructs in (**a**), as measured using EOP assays for phages Bas49, Bas52 and Bas57. Data represent the mean of at least 3 replicates and are normalized to the control samples lacking Zorya.

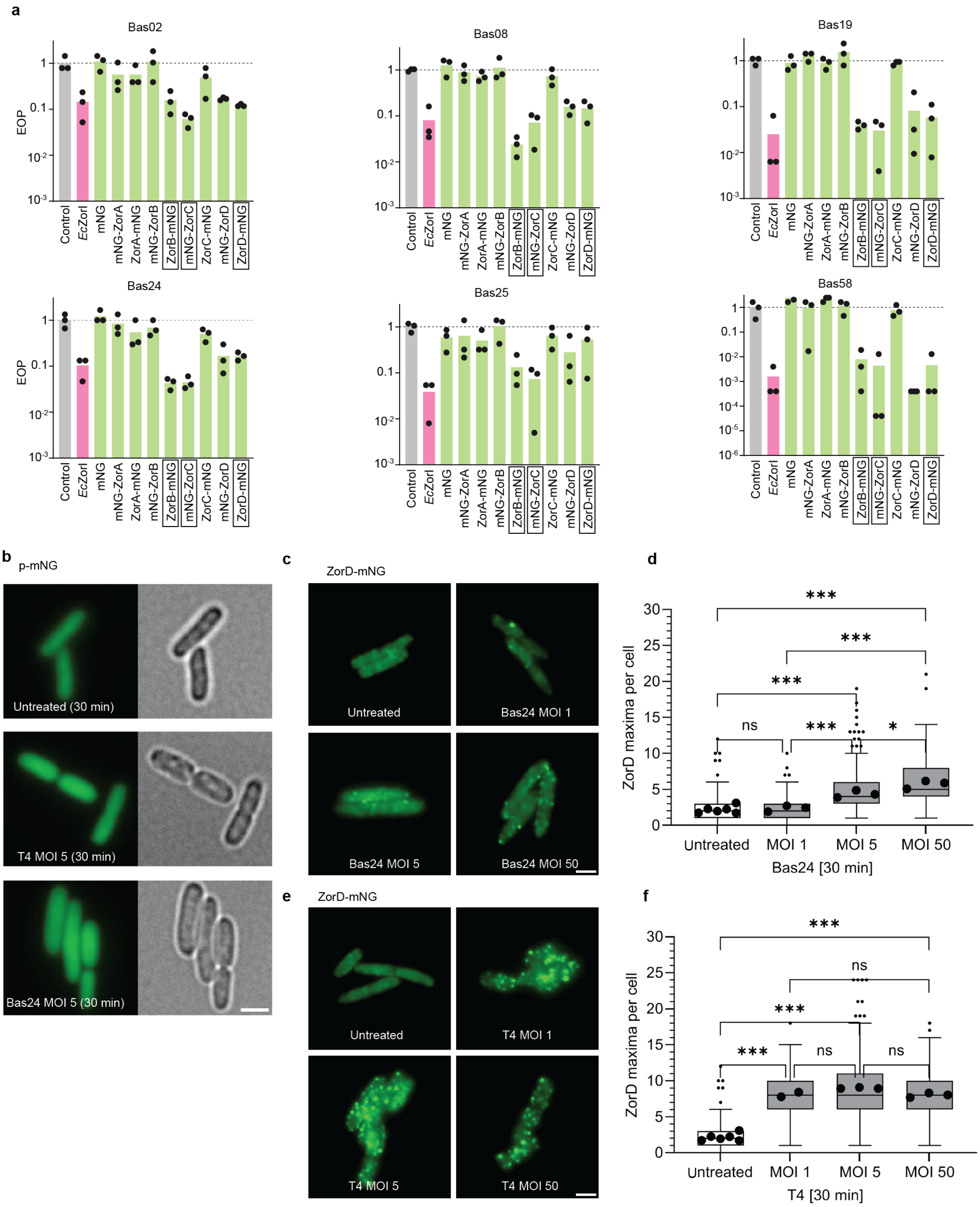

a
Bas02
Bas08
Bas19
Bas24
Bas25
Bas58
EOP
Control
EcZorI
mNG
mNG-ZorA
ZorA-mNG
mNG-ZorB
ZorB-mNG
mNG-ZorC
ZorC-mNG
mNG-ZorD
ZorD-mNG
b
p-mNG
Untreated (30 min)
T4 MOI 5 (30 min)
Bas24 MOI 5 (30 min)
c
ZorD-mNG
Untreated
Bas24 MOI 1
Bas24 MOI 5
Bas24 MOI 50
d
ZorD maxima per cell
Untreated
MOI 1
MOI 5
MOI 50
Bas24 [30 min]
e
ZorD-mNG
Untreated
T4 MOI 1
T4 MOI 5
T4 MOI 50
f
ZorD maxima per cell
Untreated
MOI 1
MOI 5
MOI 50
T4 [30 min]

**Extended Data Figure 10. ZorD recruitment during phage invasion at increasing MOIs and mNeongreen localization upon phage exposure.**

**a**, The effects of the mNeongreen (mNG) fusions to *Ec*ZorI components on anti-phage defense, as measured using EOP assays for phages Bas02, Bas08, Bas19, Bas24, Bas25 and Bas58. Data represent the mean of at least 3 replicates and are normalized to the control samples lacking *Ec*ZorI. The boxed constructs (ZorB C-terminal mNG fusion: ZorB-mNG; ZorC N-terminal mNG fusion: mNG-ZorB; ZorD C-terminal mNG fusion: ZorD-mNG) were used for subsequent microscopy experiments. **b**, Exemplary denoised TIRF and brightfield microscopy pictures of mNeongreen expression driven by the *Ec*ZorI native promoter (p-mNG) either untreated, exposed to T4 or Bas24 at an MOI of 5 for 30 min. Scale bar 2 µm. **c**, **e**, Exemplary denoised TIRF microscopy pictures of ZorD-mNG either untreated or exposed to increasing Bas24 or T4 MOIs of 1, 5, or 50 for 30 min. **d**, **f**, Statistical comparison of ZorD-mNG maxima between untreated and conditions stated in **c**, **e**. Means derive from at least three independent biological replicates. Scale bar 2 µm. Our data in **e** and **f** showed phage T4 infection did not result in a dose-dependent ZorD-mNG response.

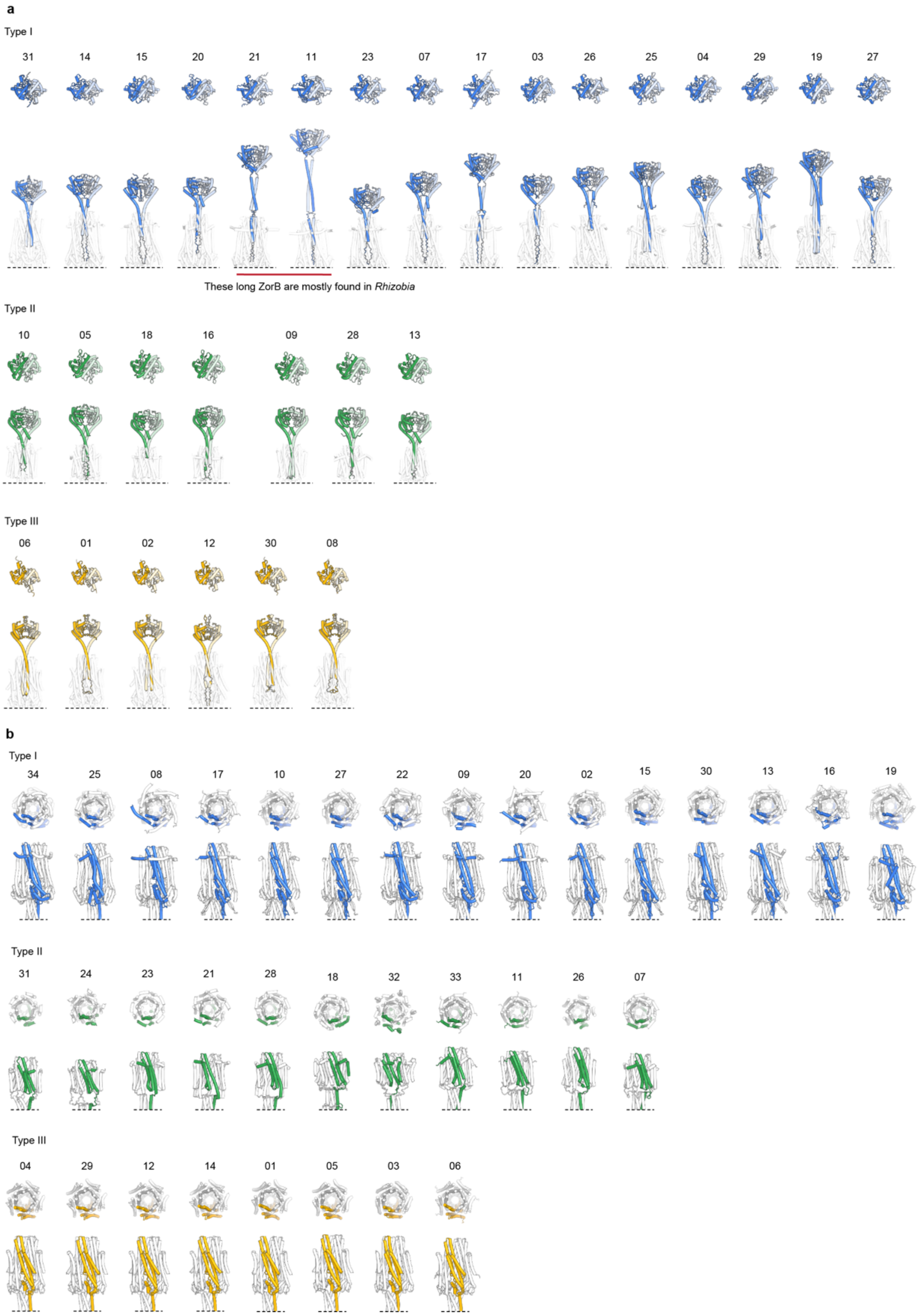


**Extended Data Figure 11. Structural prediction of the representative ZorAB complexes form different Zorya system types. a**, The predicted dimerized ZorB PGBDs. **b,** The predicted ZorAB transmembrane motor complex. One ZorA subunit is highlighted.

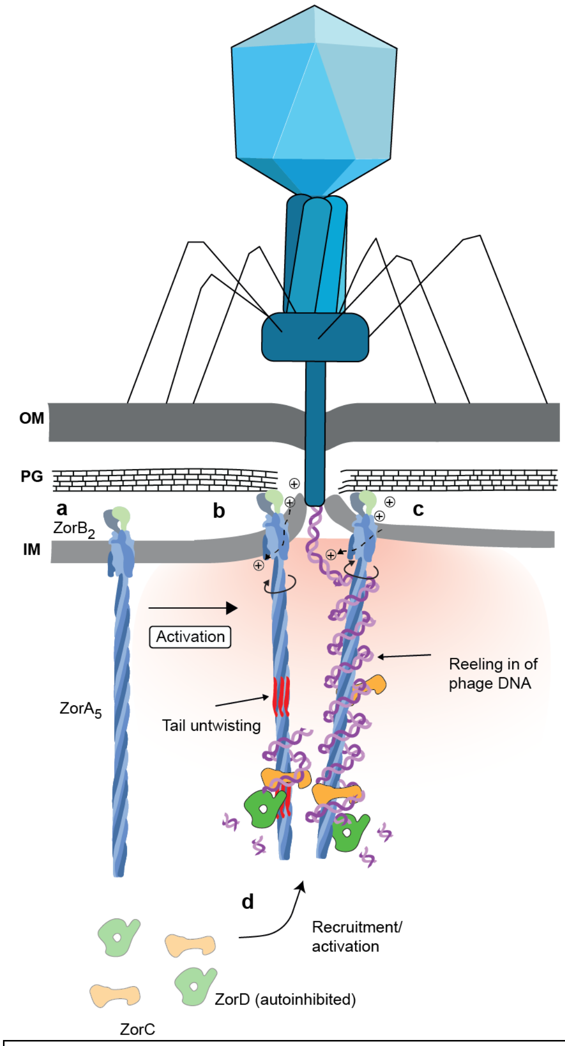


**Extended Data Fig. 12. Proposed ZorA tail untwisting and phage DNA 'reeling in' mechanism in the activated Zorya defense system.**

**a,** An inactive ZorAB embedded in the inner membrane. **b,** Phage invasion triggers ZorAB activation. The rotation of ZorA and its long intracellular tail around ZorB causes untwisting of the ZorA tail, which would recruit ZorC and ZorD**. c,** Reeling in of phage DNA around the long ZorA tail in the activated Zorya defense system. Legend Discussion: The 'reeling in' mechanism would greatly enhance phage genome localization and sequester it from interactions required for host infection. A typical double-stranded DNA phage genome is 10s of µm long, and would form a random coil with a radius of gyration similar to the size of the entire cell in the absence of constraints – thus negating any advantage of a localized nuclease defense response at the site of entry. Binding to multiple Zor complexes, perhaps via ZorC and/or D might contribute to localizing phage DNA to the entry site. Unless and until tail rotation is resisted, an almost inevitable consequence of tail rotation combined with DNA binding is that the DNA will wind around the tail like wire on a reel. If 'reeling in' of the phage DNA were an essential feature of the Zorya defense mechanism, this would explain the need for both rotation and a long ZorA tail. The length of the ZorA tail is very similar to that of a typical phage capsid into which an entire phage genome can (only just) be tightly packed, indicating that a single Zor complex may be sufficient to capture an entire genome. Rough calculations indicate that 100s of turns would be required to wind a 60 kb Bas24 genome onto a 70 nm tail, allowing the rotary ZorAB motor cumulatively to supply the necessary energy to wind and compact the phage DNA. Reeling would also tighten any loops that might form between DNA sites bound to different ZorA tails, removing any freedom for further ZorA rotation - as required by the model of activation via untwisting of the ZorA tail.
